# Separate Compartments for Chromosome Entrapment and DNA Binding during SMC translocation

**DOI:** 10.1101/495820

**Authors:** Roberto Vazquez Nunez, Laura B. Ruiz Avila, Stephan Gruber

## Abstract

Multi-subunit SMC ATPase complexes translocate on chromosomal DNA. They control chromosome structure and DNA topology, presumably by acting as DNA extrusion motors. The SMC-kleisin ring entraps chromosomal DNA. The ring lumen is strongly reduced in size by alignment of the SMC arms and upon ATP binding is divided in two by engagement of SMC head domains. Here, we provide evidence for DNA binding in the <u>S</u>MC compartment and chromosome entrapment in the <u>K</u>leisin compartment of *B. subtilis* Smc/ScpAB. We show that DNA binding at the Smc hinge is dispensable and identify an essential DNA binding site at engaged heads which faces the S compartment. Mutations interfering with DNA binding do not prevent ATP hydrolysis but block DNA translocation by Smc/ScpAB. Our findings are consistent with the notion that Smc/DNA contacts stabilize looped DNA segments in the S compartment, while the base of a chromosomal DNA loop is enclosed in the K compartment. Transfer of DNA double helices between S and K compartments may support DNA translocation.

## Introduction

Lengthwise condensation of DNA molecules into rod-shaped chromatids is a prerequisite for the faithful segregation of chromosomes during mitotic and meiotic nuclear divisions (Belmont, 2006; Kschonsak and Haering, 2015; Nasmyth, 2001). Several lines of evidence suggest that condensation relies on the formation of a series of radial loops along the chromosome axis (Earnshaw and Laemmli, 1983; Gibcus et al., 2018; Marsden and Laemmli, 1979; Naumova et al., 2013). Condensin-type SMC protein complexes are necessary for the formation of mitotic and meiotic chromosomes (Hirano, 2016; Houlard et al., 2015). *In vitro*, condensin supports ATP-dependent processive enlargement of DNA loops apparently by holding onto one DNA double helix and translocating along another, a process referred to as DNA loop extrusion (‘LE’) (Ganji et al., 2018). Presumably using a similar principle, the related cohesin complex organizes chromosomes during interphase to promote gene regulation and maybe also DNA repair and recombination (Merkenschlager and Nora, 2016; Rao et al., 2017). How exactly the highly-elongated SMC complexes use ATP binding and ATP hydrolysis for DNA translocation is an important question in chromosome biology.

Chromosome segregation in bacteria also involves SMC protein complexes (Gruber, 2011; Hirano, 2016). Chromosome conformation capture experiments indicate that MukBEF – a distantly related SMC complex found in enterobacteria – may generate a series of DNA loops covering most parts of the *Escherichia coli* chromosome (Lioy et al., 2018). The more widely-distributed Smc/ScpAB complex is thought to initiate DNA translocation mainly or exclusively from within the same chromosomal region – surrounding the replication origin on the bacterial chromosome (Gruber and Errington, 2009; Minnen et al., 2016; Sullivan et al., 2009). Rather than promoting lengthwise DNA compaction, DNA translocation by Smc/ScpAB aligns the two chromosome arms flanking the replication origin (Minnen et al., 2016; Wang et al., 2017; Wang et al., 2015). By doing so, it may help DNA topoisomerase IV to untangle nascent sister chromosomes and enable efficient chromosome segregation, particularly during fast growth under nutrient-rich conditions (Burmann and Gruber, 2015; Gruber and Errington, 2009; Gruber et al., 2014; Wang et al., 2014). Despite obvious regulatory differences, structural similarities imply that a fundamentally conserved mechanism underlies DNA translocation by Smc/ScpAB and other SMC complexes.

Smc/ScpAB is recruited to the replication origin region of the chromosome by the ParABS system in several bacteria including *Bacillus subtilis,* (Gruber and Errington, 2009; Minnen et al., 2011; Sullivan et al., 2009). Recruitment requires ParB protein bound to *parS* sequences, while the ParA ATPase is dispensable. Smc/ScpAB then translocates onto DNA flanking the *parS* loading site. All steps of this process, recruitment to and release from *parS* as well as DNA translocation, are intimately linked with the Smc ATPase cycle (Minnen et al., 2016; Wang et al., 2017; Wang et al., 2018; Wilhelm et al., 2015).

Smc proteins harbor a ~50 nm long antiparallel coiled coil ‘arm’ with a hinge dimerization domain at one end and a head domain at the other end. Two Smc proteins associate by homotypic interactions of the hinge domains and of the two Smc arms, thus also bringing the two head domains in close proximity (<u>J</u>uxtaposed heads or J heads) (Figure 1A) (Diebold-Durand et al., 2017; Hirano and Hirano, 2004; Soh et al., 2015). N-and C-terminal domains of a single ScpA subunit (kleisin) bind to opposite Smc heads, while the central region of ScpA associates with a dimer of ScpB proteins (Burmann et al., 2013; Palecek and Gruber, 2015). The pentameric Smc/ScpAB complex thus folds into a highly-elongated, rod-shaped particle. While the circumference of the tripartite SMC/kleisin ring allows for a large lumen, the alignment of the Smc arm strongly reduces its size by closing the S compartment, i.e. the lumen enclosed by Smc hinge, arms and heads (Figure 1A).

**Figure 1.**
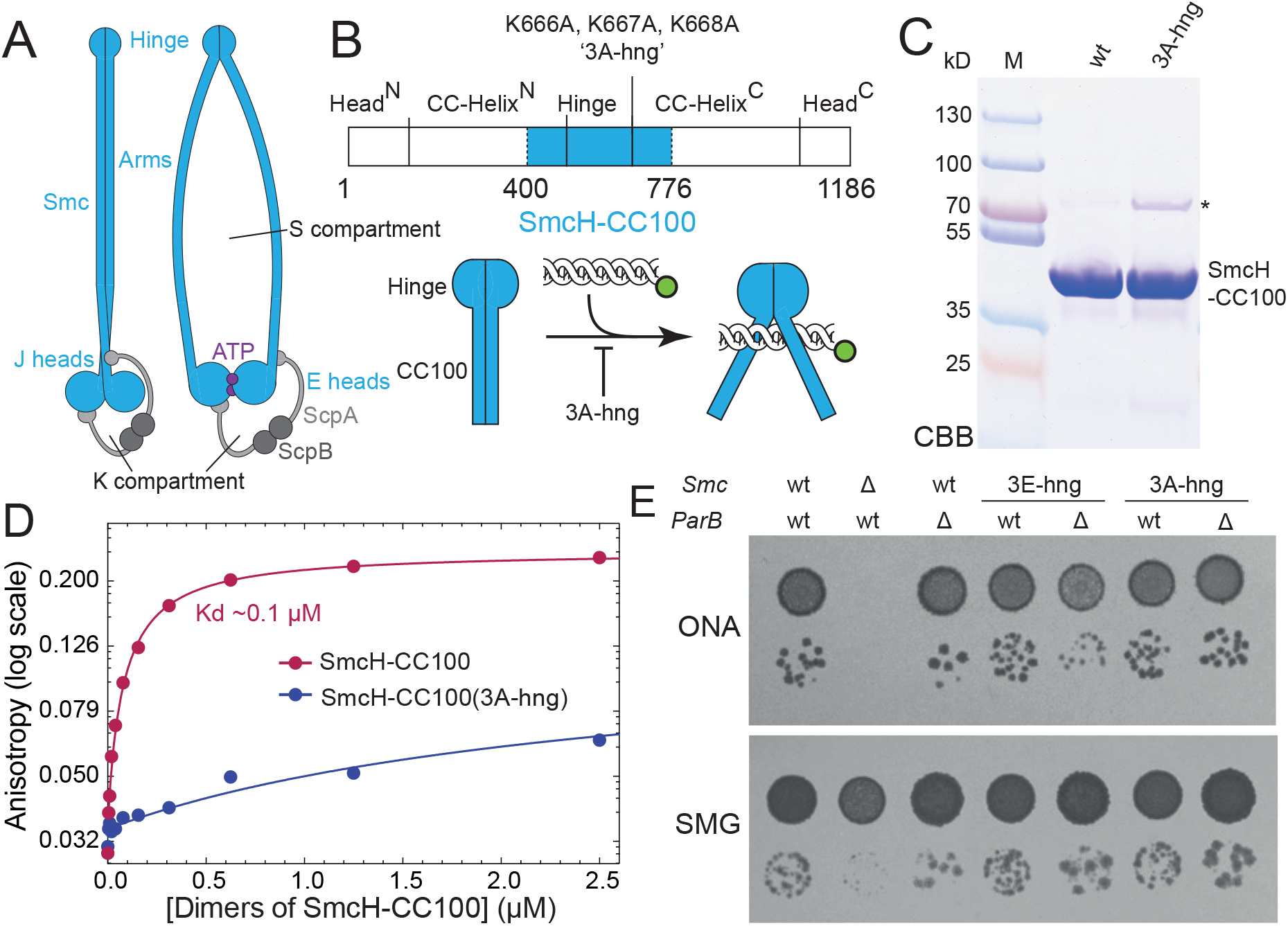
DNA binding and viability of Smc hinge mutants. (A) Schematic representation of the Smc/ScpAB complex. The J heads arrangement with aligned Smc arms is illustrated on the left panel, while a E heads state formed upon ATP binding is shown on the right. (B) Schematic view of the SmcH-CC100 construct carrying the K666A, K667A, K668A (3A-hng’) mutation. (C) Coomassie Brilliant Blue (CBB) staining of purified SmcH-CC100. The asterisk denotes an impurity present in the 3A-hng variant. Size of marker proteins (lane M) is given in kilodalton (kD). (D) DNA binding as measured by fluorescence anisotropy changes with increasing protein concentration using fluorescein-labelled 40 bp double-stranded DNA. Data points are shown as dots. The line corresponds to a non-linear regression fit of the experimental data (see experimental procedures). A representative example of three replicates is shown. (E) Colony formation by *Smc(3E-hng)* and *Smc(3A-hng)* mutant strains. Diluted overnight cultures were spotted on nutrient-rich medium (ONA) or nutrient-poor medium (SMG) and grown at 37 °C for 16 and 24 hours, respectively. See also Figure S1 and Table S2.

Like in ABC transporters, the active Smc ATPase is formed by two nucleotide binding domains (the Smc heads) and two ATP molecules (Hirano et al., 2001; Hopfner, 2016). Each ATP molecule binds to the Walker box motifs of one head and to the ABC signature motif of the other head, thus gluing the two heads together (Engaged heads or E heads) (Lammens et al., 2004). On top of E heads, the two Smc arms emerge at a distance from one another and at an open angle (Figure 1A) (Diebold-Durand et al., 2017; Kamada et al., 2017). Head engagement thus opens the S compartment (at least partially) and at the same time separates it from the K compartment, which is enclosed by kleisin ScpA and the Smc heads). Disengagement of E heads upon ATP hydrolysis merges S and K compartments into a single lumen enclosed by the SMC-kleisin ring. When the two Smc arms re-align, the lumen of the SMC-kleisin ring is reduced and converted into the K compartment (Diebold-Durand et al., 2017). Whether and how the opening, closure, fusion or fission of compartments is related to DNA translocation is poorly understood. Conceivably, it may support DNA translocation by passing DNA from one compartment to another bearing resemblance with ligand transport by ABC transporters or DNA strand passage by DNA topoisomerase II enzymes (Hopfner, 2016; Vos et al., 2011).

Here, we map locations for physical DNA binding and DNA entrapment in *B. subtilis* Smc/ScpAB. We identify a novel DNA binding site located at the Smc heads being accessible only from within the S compartment. The DNA binding site is formed on top of E heads and is essential for Smc function. The DNA interface appears conserved in highly diverged SMC-like proteins (Liu et al., 2016; Seifert et al., 2016; Woo et al., 2009) and may be a defining feature of the large family of chromosomal ABC ATPases. DNA binding is essential for Smc DNA translocation but has only limited impact on the ATP hydrolysis rate. In contrast, a previously identified DNA binding site at the hinge – capable of more strongly stimulating the Smc ATPase – is dispensable for Smc function and DNA translocation. We detect DNA entrapment within the K compartment but not within the S compartment, a feature which appears conserved in yeast cohesin (Chapard et al.). Conceivably, the S compartment binds to a strongly bent DNA segment. DNA transactions involving DNA transfer between S and K compartments may form a basis for SMC DNA translocation and DNA loop extrusion.

## Results

### A high-affinity DNA binding site at the Smc hinge is dispensable for Smc function

DNA binding capabilities have been reported for hinge domains of several SMC complexes (Alt et al., 2017; Arumugam et al., 2003; Chiu et al., 2004; Griese and Hopfner, 2011; Griese et al., 2010; Hirano and Hirano, 2006; Soh et al., 2015; Uchiyama et al., 2015). In *B. subtilis,* the substitution of three consecutive lysine residues (K666, K667 and K668) at the hinge/coiled coil junction for glutamate strongly reduces DNA binding and DNA-stimulation of ATP hydrolysis in purified preparations of Smc (Hirano and Hirano, 2006). The physiological importance of the three lysine residues, however, has remained elusive so far. Since the triple glutamate mutation (here denoted as ‘3E-hng’ for ‘three glutamates in hinge’) was not tolerated well in recombinantly expressed fragments of the Smc protein, we also generated a corresponding triple alanine mutant (denoted as ‘3A-hng’). First, we introduced the 3A-hng mutation into the SmcH-CC100 construct harboring the Smc hinge and 100 residues of the adjacent coiled coil (Soh et al., 2015) (Figure 1B). We purified wild-type and 3A-hng SmcH-CC100 to measure binding to 40 bp fluorescein-labelled dsDNA by fluorescence anisotropy (Figure 1B-C, S1). Robust DNA binding by wild-type SmcH-CC100 was observed, as reported previously (Soh et al., 2015). In contrast, the 3A-hng mutant exhibited strongly reduced DNA binding even at elevated protein concentrations (Figure 1D).

Next, we introduced *Smc(3A-hng)* into the endogenous *Smc* locus by allelic replacement. Cells carrying *Smc(3A-hng)* as sole source of Smc protein efficiently formed colonies on nutrient-rich medium (‘ONA’) at 37 °C, demonstrating normal or near-normal Smc function despite being defective in DNA binding at the hinge (Figure 1E). To sensitize for partial loss of Smc activity, we then deleted the gene encoding for the Smc loader ParB (Gruber and Errington, 2009; Wang et al., 2018). We found that also in the absence of ParB protein, Smc(3A-hng) supported growth on nutrient-rich medium surprisingly well (Figure 1E). Similar results were obtained with Smc(3E-hng), albeit colonies grew slightly slower in a *parB* deletion background (Figure S1). Six lysine residues on the dimer of Smc hinge domains can thus be substituted for six glutamate residues without major consequences for Smc function. The positively charged surface at the Smc hinge is dispensable for Smc function in *B. subtilis.* Moreover, the Smc hinge domain can be replaced altogether by the structurally unrelated Rad50 Zinc hook (Burmann et al., 2017). In the light of these results, a reassessment of the physiological relevance of hinge-DNA binding in other SMC complexes seems warranted.

### Identification of surface-exposed Smc head residues required for Smc function

The results presented above imply that Smc/ScpAB harbors one or several additional sites for DNA binding to support its putative DNA motor function. In case of MukB and the SMC-like protein Rad50, DNA binding has been demonstrated at the opposite end of the long coiled-coil arm. On top of MukB and Rad50 E heads, a DNA binding interface is formed by both heads contributing positively charged residues for DNA binding (Liu et al., 2016; Seifert et al., 2016; Woo et al., 2009). Reverting the charge of such residues in yeast Rad50 protein renders cells sensitive to DNA damaging agents – similar to Rad50 deletion mutants – implying that DNA binding to Rad50 heads is important for the DNA repair function of the Rad50-Mre11 complex (Seifert et al., 2016).

To assess whether such a DNA binding interface may exist in SMC proteins, we selected conserved *Bs* Smc head residues with surface-exposed positively-charged side chains (Figure 2A). We mutated five lysine and three arginine residues individually to glutamate. The mutant strains showed wild-type-like growth on nutrient rich medium (Figure S2A). However, five of the eight mutations could not be combined with *parB* deletion indicating that their charge-reversal compromises Smc function by potentially hindering DNA binding. To elucidate whether these five residues fulfill largely redundant or additive functions, we decided to systematically test combinations of mutations. Before doing so, we changed from glutamate to alanine substitutions aiming to reduce any undesired impact on protein folding and stability. As expected, the five single alanine mutants did not display obvious growth phenotypes (Figure S2B). Likewise, the ten double-alanine mutants grew robustly on nutrient rich medium (Figure S2C). However, when we combined three or more mutations, we identified alleles with strong growth phenotypes (Figure 2B-C). There is no obvious pattern between the degree of growth inhibition and the nature of the underlying mutations. We suspect that all five positively-charged residues contribute to Smc function and that some residue combinations perform better than others. We selected mutant K60A, R120A and K122A (denoted as ‘3A-hd’ for ‘three alanines in head’) for further characterization. Amongst the ten triple-alanine mutants, 3A-hd showed a particularly severe growth phenotype. However, 3A-hd was not as severely impaired in growth as the quintuple mutant, possibly due to residual DNA binding in 3A-hd (Figure 2C). Importantly, all tested triple-, quadruple- and quintuple-alanine mutant proteins were expressed at similar levels as the wild-type protein (Figure 2B-C).

**Figure 2.**
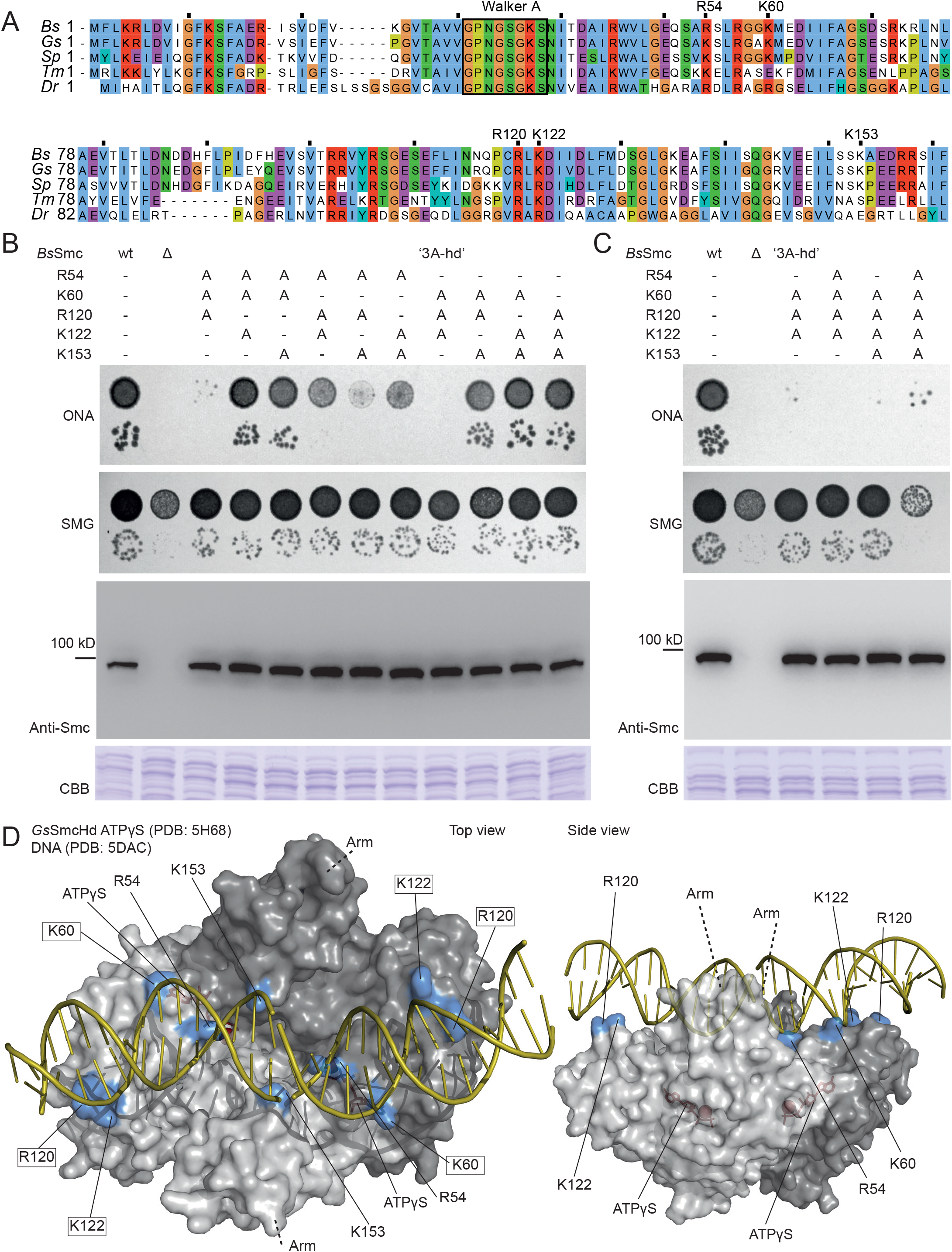
Identification of surface-exposed Smc head residues required for Smc function. (A) Multiple sequence alignment of the N-terminus of five bacterial Smc proteins (*Bs, Bacillus subtilis; Gs, Geobacillus stearothermophilus; Sp, Streptococcus pneumoniae; Tm, Thermotoga maritima; Dr, Deinococcus radiodurans*). The conserved Walker A box motif is indicated for reference. Residues chosen for further analysis are denoted. (B) Characterization of *Smc* alleles harboring triple alanine mutations in the hinge. Top panel: Colony formation by dilution spotting as in Figure 1E. Bottom panels: Cellular expression levels of Smc variants determined by immunoblotting with polyclonal antibodies raised against full-length Smc protein. Coomassie Brilliant Blue (CBB) staining of cell extracts on separate gels is shown as control for uniform protein extraction. (C) *Smc* alleles harboring quadruple and quintuple alanine mutations in the head. As described in (B). (D) Surface representation of the structure of the *G. stearothermophilus* SmcHd-ATPγS complex (PDB:5H68) (in gray colors) superimposed onto Rad50Hd-ATPγS-DNA (PDB:5DAC) (only DNA is shown – in yellow colors) (Kamada et al., 2017; Seifert et al., 2016). The side chains of putative DNA binding residues are marked in blue colors. ATPγS is shown in stick representation in red colors. Residues mutated in 3A-hd are marked by boxes (left panel only). Top view (left panel) and side view (right panel). See also Figure S2

To explore whether the five identified residues are appropriately arranged on the Smc head to jointly promote dsDNA binding, we aligned the structure of *Geobacillus stearothermophilus* ATPγS E heads (PDB: 5H68) with two available Rad50-Mre11-DNA co-crystal structures (PDB: 5DAC and 5DNY) (Kamada et al., 2017; Liu et al., 2016; Seifert et al., 2016). In the superimpositions (RMSD 3.1 and 2.3 Å, respectively) the identified positively charged residues are in proximity of the negatively charged phosphate backbone of DNA (Figure 2D and S2). Conceivably, an essential DNA binding interface exists on top of E heads Smc.

### Isolated Smc heads associate with DNA

To test whether Smc proteins indeed associate with DNA via the head domain, we co-expressed *B. subtilis* Smc heads (‘SmcHd’) with a His-tagged N-terminal fragment of ScpA (‘ScpA-N’) (Figure 3A). The latter provides a tag for affinity purification and stabilizes recombinant SmcHd preparations (Burmann et al., 2013). The 3A-hd mutation did not adversely affect expression and purification of wild-type SmcHd protein or the ATP hydrolysis deficient variant (‘EQ’ for Walker B mutation E1118Q) (Figure 3B). We then measured ATP dependent dimerization of SmcHd(EQ)/ScpA-N using analytical gel filtration. In absence of ATP, wild-type and 3A-hd proteins eluted as a single peak at the volume expected for the monomeric complex (Figure 3C, S3). In presence of ATP, an additional faster eluting species was detected presumably corresponding to the ATP-induced dimer form. The dimer peak was slightly more pronounced in the 3A-hd mutant than in wild type. The positive surface charged surface on top of Smc heads may therefore decrease the efficiency of head engagement somewhat.

**Figure 3.**
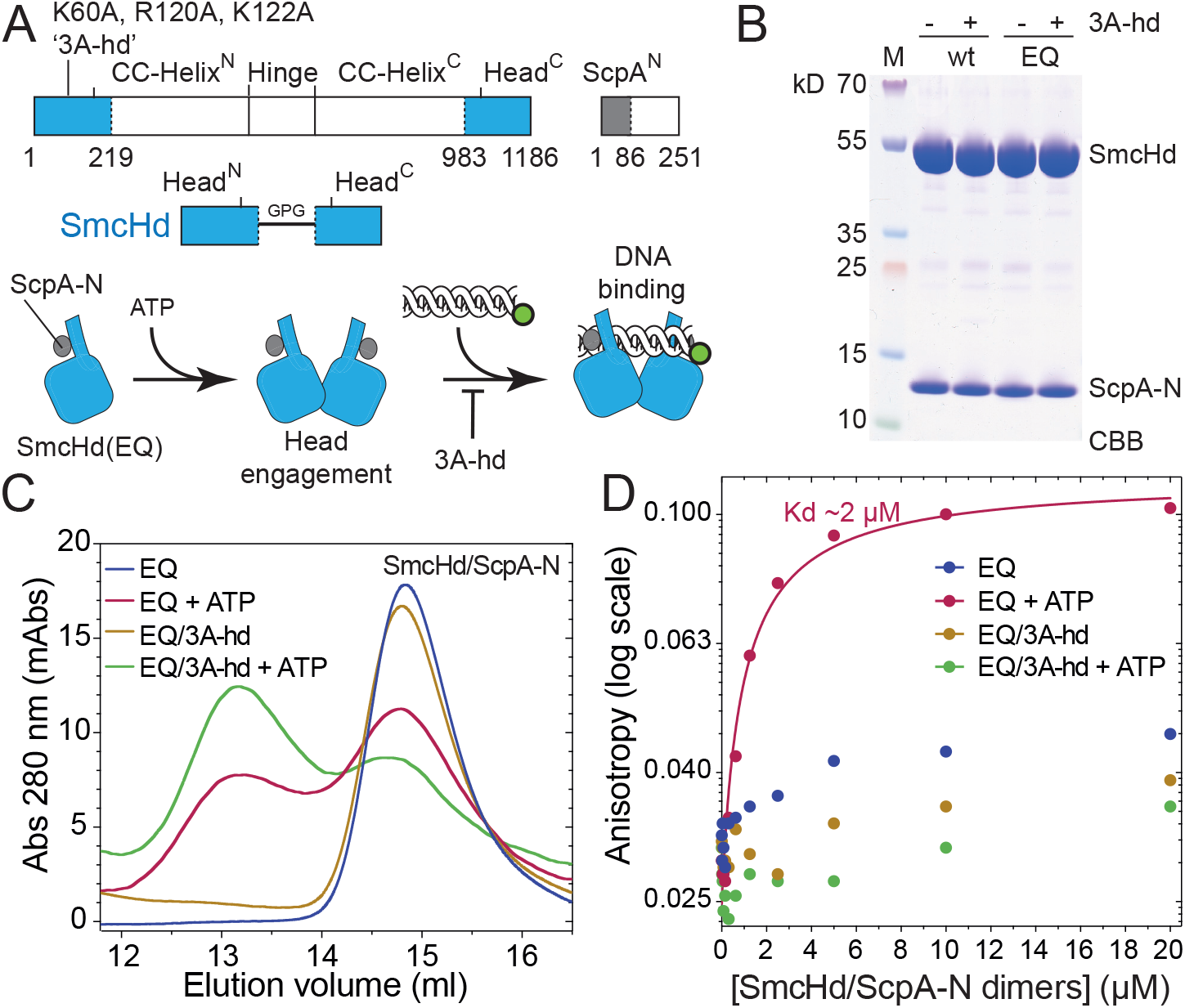
DNA binding by isolated Smc heads. (A) Construction of SmcHd/ScpA^N^-His6 with and without the ‘3A-hd’ mutations (K60A, R120A, K122A). (B) Coomassie Brilliant Blue (CBB) staining of purified preparations of SmcHd/ScpA^N^ constructs. (C) Gel filtration profiles of SmcHd/ScpA^N^ preparations eluted from a Superdex 200 10/300 column. Samples with ATP were run in buffer supplemented with 1 mM ATP. EQ denotes the Walker B ATP hydrolysis mutation (E1118Q) in SmcHd. (D) DNA binding curves from fluorescence anisotropy titrations as in Figure 1D. See also Figure S2 and Table S2.

Next, we investigated DNA binding by fluorescence anisotropy as described above (Figure 3D, S3). SmcHd(EQ)/ScpA-N associated poorly with 40 bp dsDNA in the absence of ATP but showed robust DNA binding when ATP was added. The 3A-hd variant, however, failed to exhibit detectable DNA binding either in the absence or presence of ATP. We conclude that engaged Smc heads bind DNA and that alanine mutations on top of Smc heads abolish DNA binding but do not negatively impact Smc head engagement.

### The Smc ATPase is controlled by two distant DNA binding sites in the S compartment

ATP hydrolysis is required for Smc function in *B. subtilis* (Minnen et al., 2016; Wang et al., 2018). The lethality of Smc(3A-hd) might thus be explained by a pronounced defect in ATP hydrolysis. To address this possibility, we measured ATP hydrolysis using an enzyme coupled assay, first in the purified preparations of SmcHd/ScpA-N. The 3A-hd mutant displayed mildly increased – rather than reduced – ATP hydrolysis rate when compared to wild-type (Figure 4A). This increase is likely explained by the elevated level of E heads in SmcHd(3A-hd)/ScpA-N (Figure 3C). Of note, we observed a quadratic change in the ATP hydrolysis rate (*V_max_*) with increasing SmcHd concentrations indicating that protein dimerization is the rate-limiting step in ATP hydrolysis by SmcHd (but not full-length Smc) (Figure S4).

**Figure 4.**
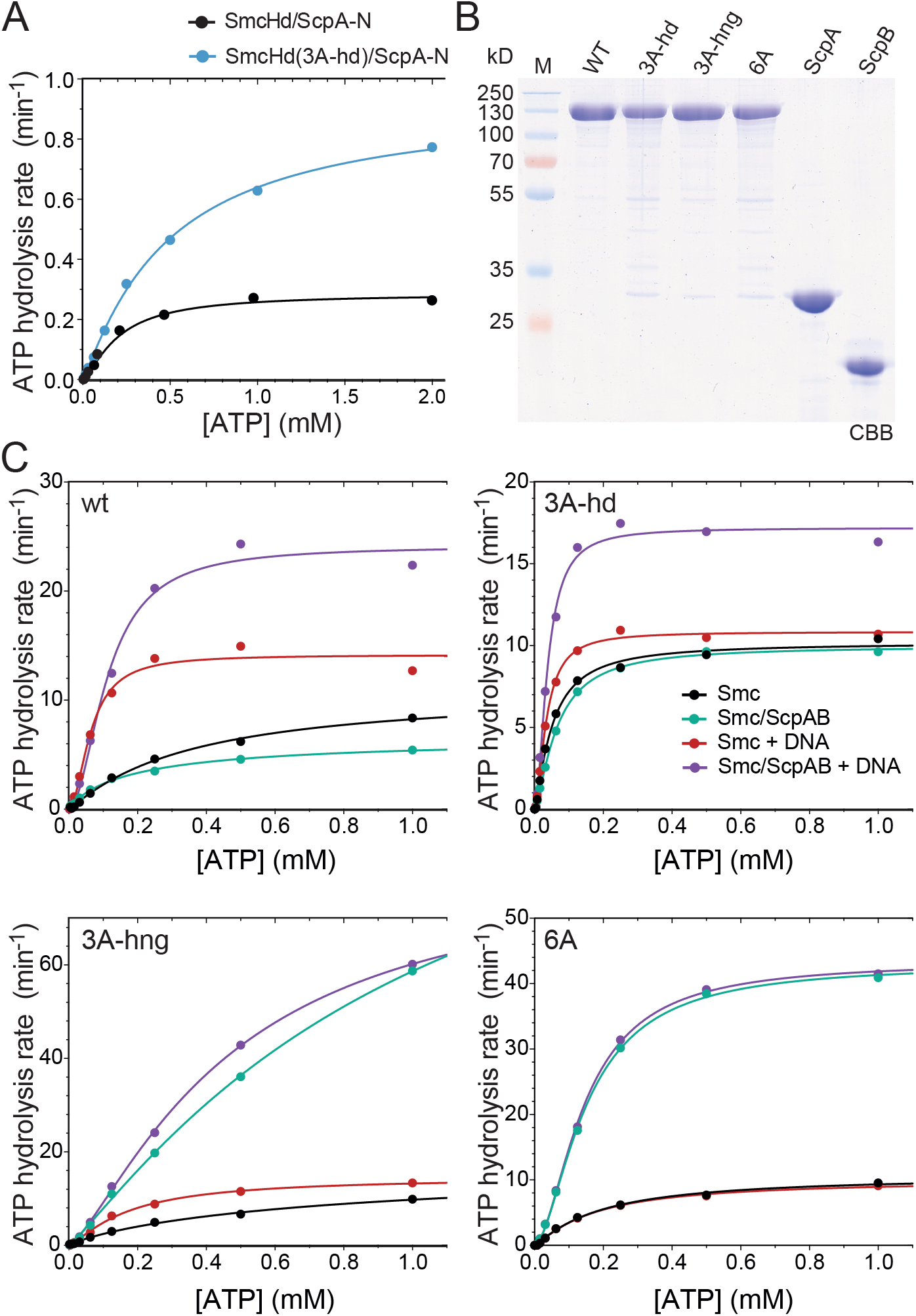
DNA stimulation of ATP hydrolysis by Smc heads and Smc/ScpAB holo-complexes. (A) ATP hydrolysis rate of SmcHd/ScpA^N^ at 10 μM protein concentration at increasing concentrations of ATP as measured by an enzyme-coupled assay. The line represents a non-linear regression fit of the Hill equation (see Experimental procedures). A representative replicate of three independent measurements is shown. (B) Coomassie Briliant Blue (CBB) staining of purified preparations of untagged, full-length Smc, ScpA and ScpB proteins. (C) ATP hydrolysis rates of full-length Smc proteins and holo-complexes at 0.15 μM Smc dimer concentration and equivalent concentratons of ScpB dimers and ScpA. As in (A) using proteins shown in (B). See also Figure S4, and Tables S3 and S4.

Next, we measured ATP hydrolysis in a more physiological setting using full-length Smc in the absence and presence of ScpAB and DNA (Figure 4B). Full-length *B. subtilis* Smc dimers exhibited a significantly higher basal ATPase activity than isolated Smc heads even at high protein concentrations (Figure 4A, C, S4). This activity was further stimulated by DNA and even more by DNA and ScpAB.

Surprisingly, the 3A-hd mutation had only limited impact on the Smc ATP hydrolase rate: While basal activity was slightly increased, and DNA stimulation slightly reduced, ATP hydrolysis rates in the presence of DNA and ScpAB were comparable to wild-type Smc under the same conditions. Since Smc mutants with much stronger defects in ATP hydrolysis display rather mild growth phenotypes (Wang et al., 2018), the loss of Smc ATPase activity or its deregulation is an unlikely cause for the lethality of 3A-hd. DNA binding on top of E heads must have a crucial role in Smc function apart from regulating the Smc ATPase.

As expected, the Smc(3A-hd) ATPase showed residual DNA stimulation, which presumably arises from DNA binding at the Smc hinge. To determine whether this is indeed the case (or whether there are other yet unidentified DNA binding sites), we combined hinge and head alanine mutations to create a sextuple alanine mutant protein. DNA stimulation of the Smc ATPase was totally abolished in the Smc(6A) protein and in Smc(6A)/ScpAB holo-complexes (Figure 4C). The Smc ATPase thus becomes insensitive to the presence of DNA when DNA binding is hindered at the hinge and at the heads, strongly implying that there are no other DNA binding sites on Smc/ScpAB capable of regulating the Smc ATPase in the absence of hinge DNA binding and head DNA binding. Importantly, the ATP hydrolysis rate of Smc(3A-hd) and Smc(6A) holo-complexes is close to wild-type levels. Defective ATP hydrolysis therefore does not explain the lethality of Smc(3A-hd).

### DNA binding by the Smc heads is required for DNA translocation

Aiming to better define the cause of lethality, we next characterized the chromosomal association and distribution of Smc(3A-hd)/ScpAB in *B. subtilis.* We were curious whether DNA binding by Smc heads is critical for the recruitment of Smc to chromosomal loading sites, for the initiation of DNA translocation or for its processivity.

First, we tested for the chromosomal association using the chromosome entrapment assay. Briefly, we use *in vivo* cysteine cross-linking to generate a covalently closed SMC/kleisin ring. Cross-linked species entrapping DNA within this ring are then co-isolated with agarose-encapsulated intact *B. subtilis* chromosomes and analyzed by Smc-HaloTag in-gel fluorescence (see also below) (Wilhelm et al., 2015; Wilhelm and Gruber, 2017). Smc(3A-hd) protein associated normally with the kleisin ScpA as judged by the cross-linking pattern observed in the input material (Figure 5A, left side). The circular species derived from Smc(3A-hd) proteins, however, were completely depleted from the chromosome plugs, similar to covalent rings obtained for ATP binding mutant Smc (Figure 5A) or for non-functional Mini-Smc proteins with artificially shortened coiled coils (Burmann et al., 2017; Wilhelm et al., 2015). This shows that the Smc(3A-hd) protein fails to stably associate with chromosomal DNA, either because it never loads onto the chromosome or because it only entraps small loops of DNA which are released during the chromosome entrapment assay.

**Figure 5.**
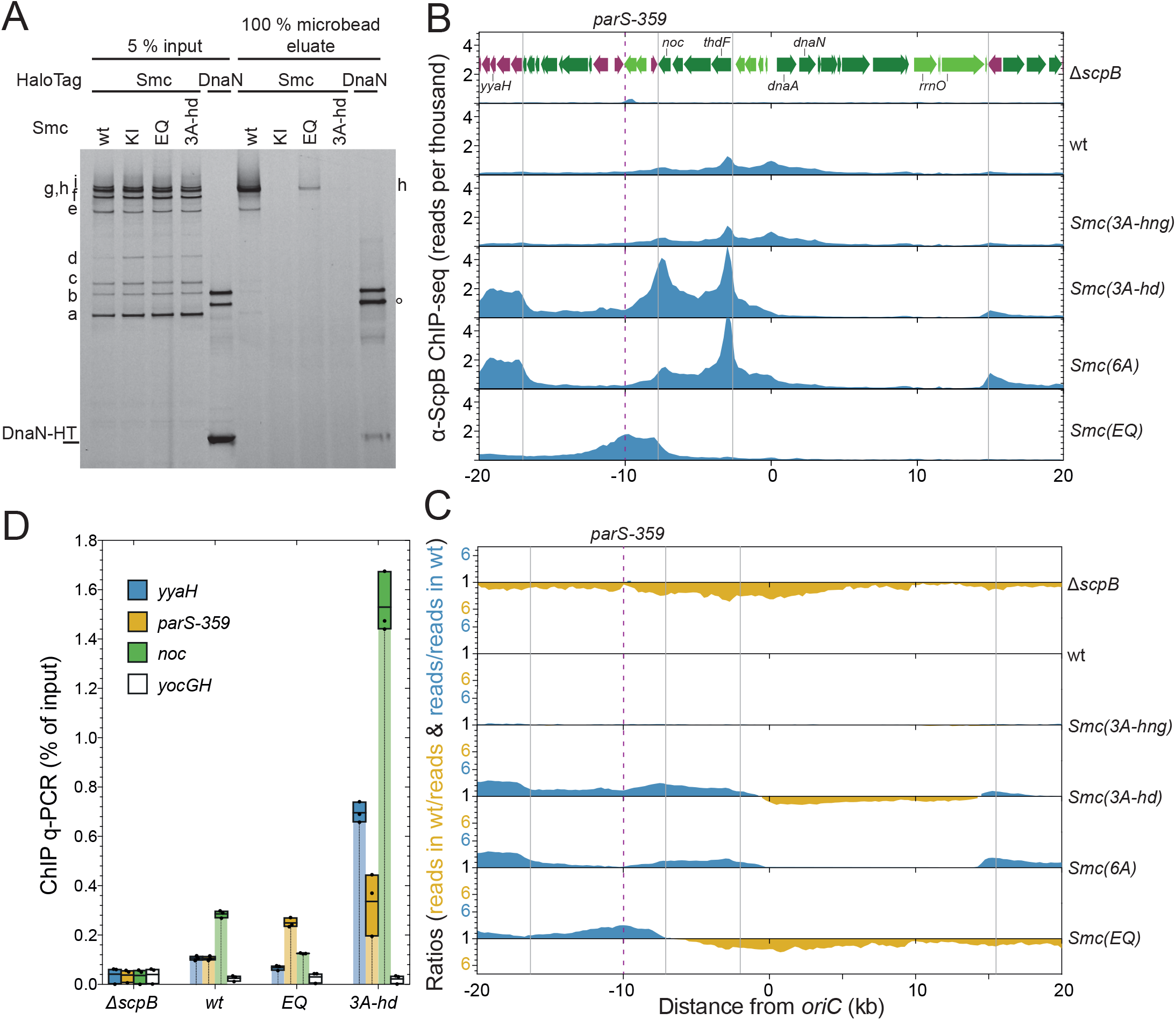
Chromosomal association and distribution of Smc(3A-hd). (A) Chromosome entrapment assay. Smc-HaloTag and DnaA-HaloTag species labelled by Halo-TMR in input and eluate fractions were analyzed by SDS-PAGE and in-gel fluorescence detection. Samples with DnaN-HaloTag were used as positive and negative controls. They were prepared by mixing cultures of DnaN-HaloTag, DnaN (N114C, V313C)-HaloTag and untagged DnaN strains at a ratio of 1:1:8. The double cross-linked DnaN-HaloTag species (denoted by °) is preferentially retained in agarose beads. All Smc-HaloTag samples harbor cysteine pairs for cross-linking of the two Smc/ScpA interfaces and the Smc hinge. KI denotes the Walker A ATP binding mutation (K37I) in Smc; EQ (E1118Q). Asterisk marks fully cross-linked circular species of Smc-HaloTag. (B) ChIP-seq profiles in the replication origin region represented as normalized reads per thousand. ChIP was performed with a polyclonal antiserum raised against the ScpB protein. Gene organization in the origin region is shown in the *ΔscpB* panel. Genes in highly-transcribed operons are displayed in green colors. Gray lines indicate the 3’ end of highly transcribed operons. The position of *parS-359* is given by a dashed line. (C) Ratiometric analysis of wild-type and mutant ChIP-Seq profiles shown in (B). For bins with read counts equal to or greater than the wild-type sample, the ratio was plotted above the genome coordinate axis (in blue colors). Otherwise the inverse ratio was plotted below the axis (yellow). (D) α-ScpB ChIP-qPCR. Selected loci in the replication origin region and at the terminus were analyzed by quantitative PCR. Each dot represents a data point from a single experiment. The solid boxes span from the lowest to the highest value obtained; the horizontal line corresponds to the mean value calculated from three independent biological replicates. See also Figure S5 and Table S5.

To address these two possibilities, we next performed chromatin immunoprecipitation with α-ScpB antiserum followed by quantitative PCR (ChIP-qPCR) or deep sequencing (ChIP-seq). The genome-wide distribution of wild-type Smc displays characteristic shallow gradients along the two chromosome arms with highest enrichment seen in the replication origin region (Figure S5A, B). The gradient is thought to arise from chromosomal loading of Smc/ScpAB at *parS* sites and subsequent active DNA translocation onto flanking DNA. Smc(EQ) accumulates at *parS* sites but is depleted from other regions of the chromosome, presumably due to a loading or translocation defect (Minnen et al., 2016) (Figure 5B-C, S5A). We observed that Smc(3A-hng) and Smc(3A-hd) proteins produced very different ChIP-Seq profiles. The Smc(3A-hng) profile was virtually identical to wild type suggesting that high-affinity DNA binding at the hinge is not required for efficient loading and translocation. The Smc(3A-hd) profile looked superficially similar to the one of Smc(EQ) but with even higher enrichment in the replication origin region and virtually complete depletion from chromosome arm regions (Figure 5B, S5A).

While Smc(EQ) peaks overlap well with ParB peaks at *parS* sites (Minnen et al., 2016), Smc(3A-hd) reached maximal enrichment at a distance from *parS.* The most prominent Smc(3A-hd) peaks were located 8 kb downstream and 2 and 6 kb upstream of the *parB* gene, which harbors a high-affinity *parS* site, called *parS-359.* Enrichment of Smc(3A-hd) directly at *parS-359* was comparatively low (Figure 5B-C). These findings suggest that Smc(3A-hd) is efficiently recruited to *parS* and released onto flanking DNA. However, Smc(3A-hd) then fails to support efficient DNA translocation, conceivably due to compromised DNA motor activity. Intriguingly, the regions displaying highest enrichment of Smc(3A-hd) were found at the 3’ end of highly-transcribed head-on gene operons (i.e. operons oriented against the presumed direction of Smc translocation) (Figure 5B). Adjacent co-directional genes (e.g. 10-20 kb upstream of *parB*), however, displayed very low levels of Smc(3A-hd) enrichment (Figure 5B-C). Thus, Smc(3A-hd)/ScpAB motors may be held back by head-on transcription complexes and pushed forward by co-directional transcription. In contrast, wild-type Smc/ScpAB efficiently bypasses head-on transcription, albeit residual accumulation at 3’ ends of genes is seen in ChIP-Seq profiles (Figure 5B).

To ensure that these observations are not artefacts caused by deep sequencing or the lack of ChIP-Seq calibration, we analyzed selected ChIP samples by qPCR. Consistent with the ChIP-Seq profiles, we found that Smc(3A-hd) is more enriched at loci adjacent to *parS-359* than at *parS-359* itself (Figure 5D). Enrichment levels of Smc(3A-hd) are substantially higher than wt and Smc(EQ) levels. These data suggest that Smc(3A-hd) – after successfully being targeted to *parS* and released onto flanking DNA – fails to support efficient DNA translocation. Holding DNA onto engaged Smc heads may thus be critical for DNA translocation by Smc/ScpAB and possibly other SMC complexes as well. These results also indicate that Smc/ScpAB complexes can translocate away from *parS* sites without being topologically associated with chromosomal DNA. This suggests that (the 3A-hd variant of) Smc/ScpAB entraps (the base of) a DNA loop rather than a single DNA double helix (designated as ‘pseudo-topological’ association) (Srinivasan et al., 2018).

### A dynamic interplay between DNA binding at hinge and heads

To our surprise, we noticed in the ChIP-Seq profiles that the Smc(6A) mutant had a slightly more normal distribution than Smc(3A-hd) (Figure 5B-C), possibly suggesting that it supports better DNA translocation. Consistent with this notion, we discovered that Smc(6A) supports substantially better growth of *B. subtilis* on nutrient-rich medium than Smc(3A-hd) (Figure 6A and S6A). Rather than aggravating the phenotype, the hinge alanine mutations partially suppress the growth defect and the aberrant chromosomal distribution caused by the head DNA binding mutant.

**Figure 6.**
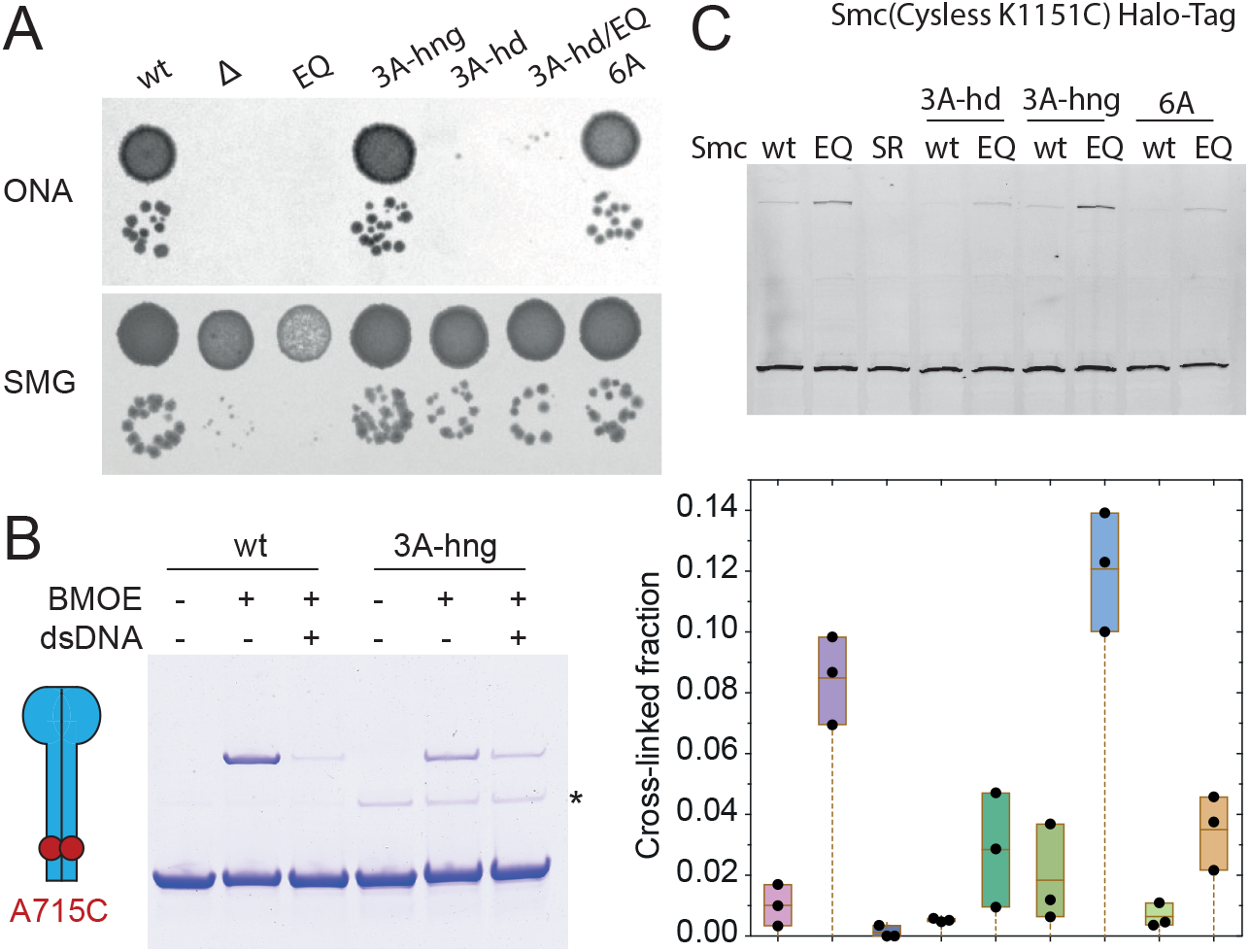
Interplay of hinge and head alanine mutations. (A) Dilution spotting of 3A-hng, 3A-hd and 6A mutants. As in Figure 1D. (B) Site-specific cysteine cross-linking of SmcH-CC100 harboring the A715C mutation in presence and absence of 40 bp dsDNA. The scheme on the left shows the positions of A715C (as red dots) in an arm-aligned conformation. Coomassie Brilliant Blue staining of protein sample with and without cross-linking (‘BMOE’). Asterisk marks an impurity; see also in Figure 1C. (C) Head engagement levels in 3A-hd, 3A-hng and 6A mutants measured by *in vivo* site-specific cross-linking using K1151C as a sensor residue. In gel-fluorescence of HaloTag-TMR ligand conjugate is shown in the top panel and the intensity quantification ratio in the bottom panel. Each dot in the graph represents an experimental data point. The boxes span from the lowest to the highest value and the horizontal line denotes the mean from three biological replicates. SR denotes the Signature motif mutation (S1090R), EQ (E1118Q). See also Figure S6.

As mentioned above, Smc(6A)/ScpAB and Smc(3A-hng)/ScpAB complexes exhibited artificially high ATPase activity in the absence of DNA (Figure 4C). The hinge alanine mutations appear to bypass the requirement for DNA binding for full ATPase activity – possibly by mimicking the DNA-bound state. Potentially related to this finding, the hinge alanine mutation influences the coiled coil architecture of SmcH-CC100 as measured by A715C cross-linking. Wild-type SmcH-CC100 gives robust cross-linking of A715C, which is strongly reduced by pre-incubation with DNA (Figure 6B) (Soh et al., 2015). The 3A-hng variant, however, displays poor cross-linking in absence (or presence) of DNA, despite forming hinge domain dimers efficiently (Figure S1). The 3A-hng protein thus preferentially adopts an ‘open-arms’ configuration (Figure 6B). We propose that DNA binding at the hinge or at the heads stabilize the same open conformation of Smc and that the 3A-hng mutation bypasses the need for DNA binding by stabilizing this conformation of Smc in the absence of DNA. Consistent with this notion, we find that 3A-hng exhibited higher levels of E Smc and E Smc(EQ) (as measured by K1151C cross-linking *in vivo*) (Minnen et al., 2016), while 3A-hd displayed decreased levels of E Smc and E Smc(EQ) (Figure 6C, S6). Importantly, we found that the lethality of the quintuple alanine head mutation (in contrast to 3A-hd) was not suppressed by 3A-hng (Figure S6). Thus, residual DNA binding at the heads seems crucial for Smc function, even in the Smc(3A-hng) mutant background.

Altogether, these findings suggest that DNA binding on top of Smc heads has essential structural roles during the Smc DNA motor cycle. It may help to dilate the S compartment as well as to promote DNA capture. Accordingly, the S compartment in Smc(3A-hd) may fail to open and/or capture DNA; in Smc(6A) however, DNA capture might be promoted by spontaneous opening of the S compartment caused byunstable alignment of the Smc arms at the hinge.

### Locating DNA double helices within Smc/ScpAB

SMC complexes are known to sterically entrap DNA molecules by the SMC-kleisin ring (Cuylen et al., 2011; Gligoris et al., 2014; Wilhelm et al., 2015). Aiming to elucidate whether DNA is entrapped within the SMC-kleisin ring when bound at the hinge or at the heads, we determined whether chromosomal DNA is located in the S compartment (or the K compartment) of Smc/ScpAB using the chromosome entrapment assay (Wilhelm et al., 2015; Wilhelm and Gruber, 2017). To do so, we covalently closed the S compartment or K compartment using cysteine cross-linking as described above.

We combined cysteine residues for cross-linking of J heads with a pair of cysteines at the Smc hinge – together locking a J-S compartment – or with cysteine pairs at both Smc/ScpA interfaces – locking a J-K compartment of Smc/ScpAB (Figure 7A). We consistently failed to co-isolate species harboring covalently enclosed S compartments with intact chromosomal DNA in agarose plugs. However, we reproducibly detected co-isolation of species with covalently enclosed K compartments. For example, the combination of Smc head S152C and Smc hinge R558C and N634C allows efficient cross-linking of Smc-HaloTag protein (Diebold-Durand et al., 2017; Wilhelm et al., 2015), but none of the cross-linked species were retained in agarose plugs (Figure 7B). Conversely, when Smc(S152C) was combined with the two Smc-ScpA cysteine pairs, a single species – in all likelihood the circular form – was reproducibly detected in the eluate sample (Figure 7B). Recovery of this J-K compartment species was consistently less efficient than the recovery of the corresponding species for the full SMC-kleisin ring, possibly indicating partial DNA occupancy in the J-K compartment, reduced levels of J Smc/ScpAB on the chromosome or technical limitations such as uneven cross-link reversal during the chromosome entrapment assay. Importantly, the retention of the covalently-closed circular species was fully dependent on ScpB protein and on ATP binding by Smc (Figure 7B). DNA entrapment by the J-K compartment is thus likely a result of a physiological chromosome loading reaction. We obtained analogous results when using cysteine pairs located at different positions along the Smc rod (for example D280C) instead of the J heads sensor residue S152C (Figure S7) (Diebold-Durand et al., 2017).

**Figure 7.**
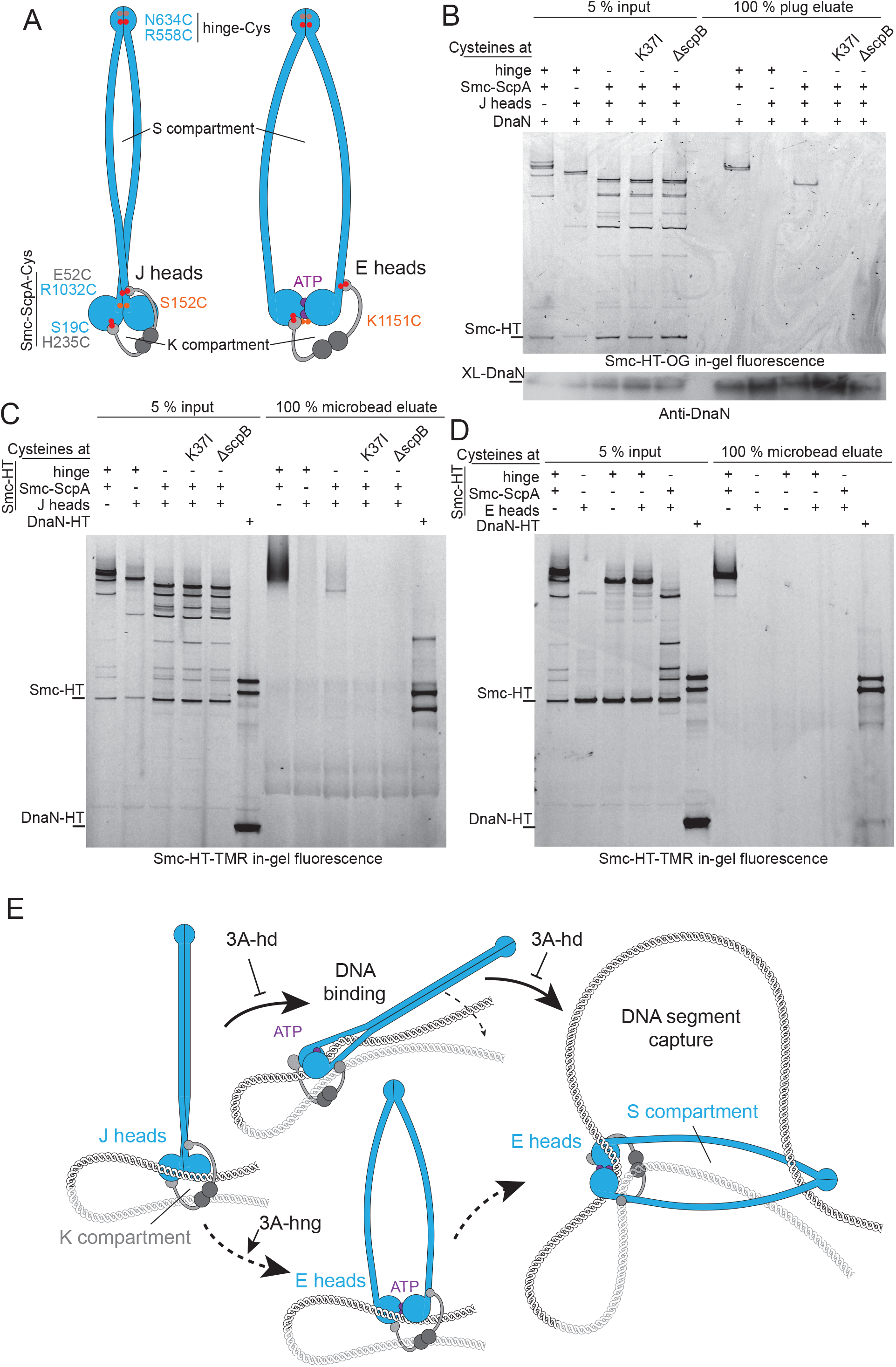
Locating DNA double helices in Smc/ScpAB. (A) Schematic view of Smc/ScpAB with J heads (left panel) and E heads (right panel). Cysteine residues previously engineered for covalent closure of the full ring compartment are indicated as dots in red colors. S152C cross-links juxtaposed heads (J heads; orange dots; left panel), while K1151C cross-links ATP engaged heads (E heads; orange dots; right panel). The Halo-Tag fusion on Smc is omitted for simplicity. (B) Chromosome entrapment assay with agarose plugs for strains harboring the J heads cysteine S152C. Cells with *Smc-HaloTag* (‘Smc-HT’) alleles were cross-linked with BMOE using cysteine pairs at the indicated protein-protein interfaces and incubated with HaloTag-OregonGreen substrate. Intact chromosomes were isolated in agarose plugs. Co-isolated proteins were analyzed by SDS-PAGE and in-gel fluorescence detection (top panel) and immunoblotting using antibodies raised against DnaN (bottom panel). K37I, Walker A ATP binding mutant of Smc. (C) Chromosome entrapment assay with agarose beads. As in (B) using agarose microbeads instead of agarose plugs and HaloTag-TMR instead of Oregon Green label. A mix of strains carrying DnaN-HaloTag (‘DnaN-HT’) with and without cysteines for cross-linking is included as positive and negative control (as in Figure 5C). (D) Chromosome entrapment assay with agarose plugs for strains harboring the E heads sensor cysteine K1151C. As in (C). (E) Speculative model for chromosome entrapment within the K compartment and DNA segment capture by the S compartment. Two DNA double helices at the base of a DNA loop are entrapped within the K compartment of J Smc/ScpAB (left panel). DNA binding to engaging Smc heads promotes opening of the S compartment and transition to full head engagement (top panel). Open S compartments capture a DNA segment and stabilize the bent DNA by physical contacts at the heads and the hinge domains (right panel). 3A-hd mutations abolish the transition to full Smc head engagement. 3A-hng mutations partially bypass the requirement for DNA binding at the heads by favoring the stochastic opening of the S compartment for random capture DNA segments (bottom panel). See also Figure S7.

Being concerned about limited sensitivity of our assay, we established a more robust version of the assay using agarose microbeads instead of agarose plugs (Figure S7A). With the improved assay, we again observed DNA entrapment in the K compartment but not in the S compartment of J Smc/ScpAB (Figure 7C). Together, these experiments demonstrate that there is J Smc/ScpAB on the chromosome. In J Smc/ScpAB, the chromosomal DNA molecule is enclosed in the K compartment while DNA seems largely or totally excluded from the S compartment – presumably by the close alignment of Smc arms in J Smc/ScpAB (Diebold-Durand et al., 2017).

Knowing that DNA binding at the heads (and possible also at the hinge) is restricted to the E heads state, we next aimed to determine the location of chromosomal DNA in E Smc/ScpAB (Figure 7D). Due to low efficiency in cross-linking of the E heads sensor K1151C (presumably caused by low frequency or duration of head engagement), these experiments are very challenging (Burmann et al., 2017; Minnen et al., 2016). We did not detect cross-linked species for the E-S compartment or E-K compartment in agarose eluates. This could be explained by insufficient cross-linking, especially in case of the K compartment that requires triple cross-linking. However, we would have expected to detect (at least the E-S compartment) species if it were efficiently entrapping DNA as double cross-linking of hinge and E heads is detectable (Diebold-Durand et al., 2017). Related experiments performed with yeast cohesin – displaying significantly higher levels E heads cross-linking – revealed entrapment of mini-chromosome DNA in the E-K compartment but not in the E-S compartment (Chapard et al.,).

Together our results describe two distinct states of Smc/ScpAB: A J state entrapping DNA within the K compartment, and an E state with open S compartment exposing two DNA binding site that possibly associates with small DNA loops. These two states are presumably fundamental intermediates of the LE process.

## Discussion

LE by SMC complexes explains many striking features of long-range chromosome organization in bacteria and in eukaryotes. Considering the wide distribution of SMC genes in bacteria and archaea, LE must have emerged early in evolution, possibly to support rapid disentanglement of fast replicating genomes. While SMC complexes have diversified and been adopted for a variety of functions in eukaryotes, the basic mechanism underlying SMC DNA translocation and LE has likely remained unaltered. The conservation of the SMC/kleisin ring is consistent with this notion. Here, we elucidate yet another putatively widely conserved feature of the tripartite core: a DNA binding site located on top of engaged Smc heads. This DNA binding site is also present in the SMC-like protein Rad50 and may be shared with other distantly related chromosomal ABC ATPases, like RecF (Liu et al., 2016; Seifert et al., 2016; Tang et al., 2018).

SMC arms have been widely considered as flexible linkers with dimensions large enough to embrace two or more 10 nm chromatin fibers. Intriguingly, we do not find evidence for DNA entrapment between the two Smc arms. Instead we find DNA located in the much smaller K compartment – at least in a significant fraction of complexes. Similar results reported for cohesin imply that this feature is widely conserved in SMC/kleisin complexes (Chapard et al.,). Together the findings support the notion that the long SMC arms serve an entirely different purpose such as enabling efficient DNA translocation. The long SMC arms may support large translocation steps needed for rapid LE or allow efficient bypass of chromosomal obstacles. Consistent with the notion of a mechanical function of Smc arms, we recently found that they tolerate changes in end-to-end distance but require a defined end-to-end orientation (Burmann et al., 2017).

Smc/ScpAB now harbors two known sites for physical contact with DNA (Figure 2, 3) (Hirano and Hirano, 2006; Soh et al., 2015). Both DNA binding interfaces are located on the Smc protein. A newly discovered DNA binding site at the heads is formed upon head engagement and is thus restricted to E Smc/ScpAB. While the exact nature of DNA binding at the hinge has remained elusive, the facts that DNA binding promotes disruption of Smc arm alignment, is inhibited by the long Smc arms and requires lysine residues located at the hinge-coiled coil junction, suggest that DNA binding occurs at the bottom of the hinge domain between the emerging Smc arms (Hirano and Hirano, 2006; Soh et al., 2015). If so, then both DNA binding sites are exposed in the E-S compartment, implying that DNA double helices engaging with either DNA binding interface are enclosed in the S compartment. However, DNA entrapment was not detected in the S compartment of Smc/ScpAB (or cohesin). DNA binding is thus either very transient or rare, or it does not lead to DNA entrapment. We favor the latter possibility and propose that a small loop of DNA occupies the S compartment with one of its ends being associated with the heads and the other with the hinge (Marko et al., 2018). In this pseudo-topological arrangement, DNA does not get interlocked with a covalently enclosed S compartment thus explaining the lack of retention during the chromosome entrapment assay (Figure 7C) (Chapard et al.). DNA loop capture has previously been reported for multiple preparations of SMC complexes and may be a widely shared feature (Kim and Loparo, 2016; Kimura et al., 1999; Kumar et al., 2017; Marko et al., 2018; Sun et al., 2013).

A surprising implication of our data is that the coordination of a single strong DNA binding site with the Smc ATPase may suffice for efficient DNA translocation. The hinge-head sextuple alanine mutant (6A) has an ATPase rate that is completely unaffected by the presence of (short double stranded) DNA molecules, supporting the notion that in the absence of the two known DNA binding sites no other hypothetical DNA contact stimulates or inhibits ATP hydrolysis. Since mutants with strongly reduced DNA binding at the hinge display essentially normal growth and apparently normal DNA translocation, DNA binding at the heads may suffice for DNA translocation. Such a scenario is difficult to reconcile with any model of DNA translocation based on walking along DNA or DNA hand-over. While the findings are largely compatible with the recently proposed DNA-segment-capture model (Diebold-Durand et al., 2017; Hassler et al., 2018; Marko et al., 2018), a simple version of the model does not provide an explanation for the difference in severity in the phenotypes of hinge and head DNA binding mutants. The model invokes an intermediate state having a DNA segment captured in the E-S compartment (Figure 7E). This intermediate is unfavorable due to entropic and energetic costs of DNA bending and needs to be stabilized by physical DNA binding at the hinge or at the heads (Marko et al., 2018). In the complete absence of physical DNA associations, DNA would rarely occupy the S compartment and DNA translocation would consequently become very inefficient (Marko et al., 2018). However, this interpretation fails to explain why head mutants have drastic phenotypic consequences while hinge mutants are tolerated well. One needs to invoke that DNA binding at the heads has additional functions apart from stabilizing a DNA intermediate in the S compartment such as fostering head engagement, opening the S compartment or competing DNA off other DNA binding sites. Our data points towards the first two possibilities (Figure 6). More generally speaking, it seems likely that an interplay between two modes of interaction, one based on physical contact with DNA and one based on steric associations of DNA in the K compartment form the mechanistic basis of DNA translocation and LE.

Our results have uncovered an additional facet of the dynamic interplay between the distantly located Smc hinge and Smc head domains (Hirano and Hirano, 2006; Minnen et al., 2016; Soh et al., 2015). We find that strong phenotypes in growth and in the chromosomal distribution of Smc caused by alanine mutations in the head are suppressed when simultaneously mutating residues in the hinge. A Smc fragment harboring the hinge alanine mutations is defective in DNA binding and it preferentially adopts an open arms conformation (Figure 6B). The suppression may be related to reduced DNA binding or facilitated opening of the Smc arms. It is in principle possible that the two DNA interfaces compete for binding to the same DNA molecule, for example in a folded conformation of SMC where hinge and head domains are brought into close proximity (Buermann et al., 2018). Weakened DNA binding at the heads, may accordingly be compensated by reduced DNA affinity at the hinge. However, our findings offer another explanation that defective alignment of the Smc arms [or defective folding of the Smc arms; (Buermann et al., 2018)] caused by the hinge alanine mutations, allows for more efficient head engagement and S compartment opening, thus bypassing a requirement for strong DNA binding at the heads (Figure 7E).

A long-standing question in the SMC field is the topological nature of SMC-DNA interactions. While sister chromatid cohesion clearly involves topological entrapment of DNA molecules by a single cohesin complex, the situation is less clear for the process of LE, by cohesin, condensin or Smc/ScpAB. A cohesin mutant, which is apparently able to translocate on DNA but unable to entrap plasmid DNA molecules, suggests that DNA translocation can occur without topological DNA entrapment (Srinivasan et al., 2018). Conversely, another recent report invokes that condensin topologically entraps DNA prior and during LE (Eeftens et al., 2017). Our data favor the former scenario, in which DNA translocation can occur without prior topological DNA entrapment. The Smc(3A-hd) mutant fails to display chromosomal DNA entrapment while it strongly accumulates on the chromosome several kilobase pairs away from the *parS* loading site, being indicative of DNA translocation. If so, then chromosomal loading of Smc/ScpAB may not require detachment of a SMC/kleisin ring interface to open the ring circumference but may equate to the capture of a first DNA segment in the lumen of the SMC/kleisin ring. This reaction may be supported by the ParB protein bound to *parS* sites. Conversion of this pseudo-topological association to a topological association in cohesin may require subsequent ring opening (Gruber et al., 2006; Murayama and Uhlmann, 2015). Alternatively, the 3A-hd mutant protein (and the beforementioned cohesin mutant) may undergo aberrant loading reaction leading to inefficient or abortive DNA translocation.

Altogether, our results provide an account of physical and steric associations between Smc/ScpAB and DNA. We identify DNA binding on top of ATP engaged Smc heads as an essential prerequisite for DNA translocation activity. All future models for DNA translocation by Smc/ScpAB (and possibly also other SMC complexes) will need to provide an adequate explanation for this requirement.

## Experimental procedures

### Bacillus subtilis strains and growth

All *B. subtilis* strains are derived from the 1A700 isolate. All phenotypes and strain numbers are listed in Table S1. Naturally competent *B. subtilis* cells were transformed as described in (Burmann et al., 2013) with longer starvation incubation time for high efficiency transformation as described in (Diebold-Durand et al., 2017). The transformants were selected on SMG-agar plates and single-colonies isolated. The strains were confirmed by PCR and sanger sequencing as required. In the case of the strains used for time course ChIP-seq, all the transformation and selection steps were done in presence of 2 mM theophylline. Dilution spot assays were done in liquid SMG at 37 °C, cells were grown to stationary phase and 9^2^ and 9^5^ -fold dilutions were spotted onto ONA (16 h incubation or as indicated) or SMG (24 h incubation) agar plates (Burmann et al., 2013).

### Protein purification

#### 6His-tagged SmcH-CC100

His6-tagged SmcH-CC100 constructs were purified following the method described in (Soh et al., 2015). The proteins were expressed from pET-28 derived vectors in *E. coli* BL21-Gold (DE3) for 24 hours at 24 °C in 1 L of autoinduction medium (OvernightExpress TB, Novagen). Extracts were sonicated in binding buffer (50 mM NaPi, pH 7.4 at 4 °C, 300 mM NaCl, 80 mM imidazole, 10 % glycerol, 1 mM DTT) and bound to Ni^2+^ Sepharose HP columns (GE Healthcare). Columns were washed with 10 column volumes (CV) of binding buffer, followed by 5 CV of 50 mM NaPi, pH 7.4 / 4 °C, 1 M NaCl, 10 % glycerol, 1 mM DTT, followed by 5 CV of binding buffer. Protein elution was done with 500 mM imidazole, pH 7.4 / 4 °C, 300 mM NaCl, 10 % glycerol, 1 mM DTT. The elution fractions were further purified on a Hi-Load Superdex 200 16/600 pg (GE Healthcare) equilibrated in 50 mM Tris-HCl, 200 mM NaCl, 1 mM TCEP, pH 7.4. Peak fractions were concentrated in Vivaspin 15 10K MWCO filters (Sartorius) and aliquots were flash frozen in liquid N2 and stored at -80 °C.

#### His6-ScpA^N^/SmcHd

The complex was purified as described in (Burmann et al., 2013). SmcHd constructs containing residues 1-219 and 983-1186 connected by a GPG linker and His6-ScpA^N^ (residues 1-86) were co-expressed in 1 L of autoinduction medium for 24 h at 24 °C. Protein extracts were prepared by sonication in binding buffer (50 mM NaPi, pH 7.4 at 4 °C with 300 mM NaCl, 10 % glycerol, protease inhibitor cocktail (Sigma) and 40 mM imidazole). The extracts were loaded onto HisTrap columns (GE Healthcare), washed with 10 column volumes of binding buffer and eluted in a gradient up to 500 mM imidazole (pH 7.4 at 4 °C) in binding buffer. Protein complexes were further purified by gel filtration on a Hi-Load Superdex 200 16/600 pg (GE Healthcare) equilibrated in 25 mM Tris-HCl, 200 mM NaCl. Finally, the samples were concentrated, flash frozen and stored at -80 °C.

#### Full-length *B. subtilis* Smc

Native Smc proteins were purified as described in (Burmann et al., 2017). Proteins were expressed from pET-22 or pET-28 derived plasmids in E. coli BL21-Gold(DE3) using ZYM-5052 autoinduction medium (Studier, 2005) during 23 h at 24°C. Cells were resuspended in lysis buffer (50 mM Tris-HCl pH 7.5, 150 mM NaCl, 1 mM EDTA, 1 mM DTT, 10% sucrose) supplemented with protease inhibitor cocktail (PIC) (Sigma) and sonicated. The soluble phase was loaded on 2 HiTrap Blue HP 5 mL columns connected in series (GE Healthcare) and eluted with a linear gradient of lysis buffer containing 1 M NaCl. The main peak elution fractions where diluted in buffer (50 mM Tris-HCl pH 7.5, 1 mM EDTA, 1 mM DTT) to a conductivity equivalent of 50 mM NaCl (≈ 8 mS/cm). The sample was loaded on a HiTrap Heparin HP 5 mL column (GE Healthcare) and was eluted with a linear gradient of buffer containing 2 M NaCl. Except for the 3Ahi/3Ahd mutant, that was loaded onto a HiTrapQ HP column (GE Healtcare) and eluted in the same conditions at the other constructs. The main peak fractions were pooled and concentrated to 5 mL in Vivaspin 15 10K MWCO filters (Sartorius). The samples were further purified by gel filtration on a XK 16/70 Superose 6 PG column (GE Healthcare) in 50 mM Tris-HCl pH 7.5, 100 mM NaCl, 1 mM EDTA, 1 mM DTT. Main peak fractions where pooled, concentrated and stored at -80°C. Protein concentration was determined by absorbance using theoretical molecular weight and molar absorption values.

#### ScpA

Native ScpA was purified using a modified protocol reported in (Fuentes-Perez et al., 2012)(Fuentes-Perez et al., 2012). The protein was expressed from a pET28 derived plasmid, transformed into E. coli BL21-Gold (DE3), cultivated in ZYM-5052 auto induction medium (Studier, 2005) at 16 °C during 28 h. Cells were harvested and resuspended in lysis buffer (50 mM Tris-HCl pH 7.5, 200 mM NaCl, 5 % glycerol) supplemented with protease inhibitor cocktail (Sigma) and sonicated. The soluble phase was loaded onto a 5 mL HiTrapQ ion exchange column (GE Healthcare) and eluted with a gradient up to 2 M NaCl. The peak fractions were resuspended in 4 M NaCl buffer up to 3 M NaCl final concentration. The sample was injected into a HiTrap Butyl HP column and eluted in a reverse gradient to 50 mM NaCl. Peak fractions were pooled and concentrated to 5 mL in Vivaspin 15 10K MWCO filters (Sartorius) and subsequently purified by gel filtration in a Hi Load 16/600 Superdex 75 pg column (GE Healthcare), equilibrated in 20 mM Tris-HCl pH 7.5, 200 mM NaCl. Protein was concentrated, aliquoted, flash frozen and stored at -80 °C.

#### ScpB

Native ScpB was purified using a protocol based on the reported in (Fuentes-Perez et al., 2012). E. coli cells harboring a pET22 derived plasmid with the coding sequence of ScpB were grown in ZYM-5052 autoinduction medium (Studier, 2005) at 24 °C during 23 h. Cells were harvested and resuspended in lysis buffer (50 mM Tris-HCl pH 7.5, 150 mM NaCl, 1 mM EDTA, 1 mM DTT) supplemented with PIC (Sigma). The material was sonicated, and the soluble phase was diluted to 50 mM NaCl, loaded onto a 5 mL HiTrap Q HP column (GE Healthcare) and eluted with a gradient up to 2 M NaCl. The sample was diluted in 4 M NaCl buffer up to a final concentration of 3 M NaCl. The sample was loaded on two 5 mL HiTrap Butyl column (GE Healthcare) connected in series. The protein was eluted with a reverse gradient down to 50 mM NaCl. The peak fractions were concentrated and purified by gel filtration using a Hi Load 16/600 superdex 200 pg equilibrated in 50 mM Tris-HCl pH 7.5, 100 mM NaCl. The fractions containing the protein were concentrated, aliquoted, flash frozen and stored at -80 °C.

#### Fluorescence anisotropy measurements

Fluorescence anisotropy was measured as reported in (Soh et al., 2015) using a 40bp DNA substrate of random sequence (TTAGTTGTTCGTAGTGCTCG TCTGGCTCTGGATTACCCGC) modified with fluorescein. The data was acquired using a Bio Tech neo plate reader with the appropriate filters at 25 °C. All measurements were done in 50 mM Tris-HCl pH 7.5, 50 mM NaCl, 2 mM MgCl2 with or without 1 mM ATP. Anisotropy values where obtained directly from the measurement software, exported and fit in GraphPad Prism 7 with the following equation:

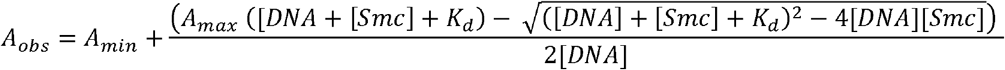

Where *A_obs_* is the experimentally measured anisotropy, *A_min_* and *A_max_* are the minimal and maximal anisotropy values respectively, *[DNA]* is de dsDNA concentration fixed at 50 nM in all our measurements, *[Smc]* is the variable Smc protein concentration and *K_d_* is the dissociation constant.

#### ATPase assay

ATPase activity measurements were done following the method of pyruvate kinase/lactate dehydrogenase coupled reaction as described in (Burmann et al., 2017). The steady state was monitored during 1 h by measuring the NADH absorbance change in a Synergy Neo Hybrid Multi-Mode Microplate reader (BioTek). The absorbance values at 340 nm were exported and fit to a straight-line equation. The slope values were transformed to rate values using the molar absorption coefficient of NADH. The rates were expressed into absolute values by correcting for the protein concentration. Data was fit to the Hill equation:

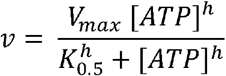

Where *v* is the ATP hydrolysis rate,*V_max_* is the maximal rate, [*ATP*] is the variable ATP concentration, *h* is the Hill coefficient and *K_0.5_* is the semi-saturation concentration.

For the protein dependence experiments (Fig S4A), in the wt protein we fit the rate dependence on protein concentration to the first order rate equation for *k_cat_*:

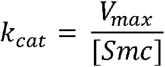

For the SmcHd/N-ScpA constructs we fit the data to the second order rate equation for *k_cat_*:

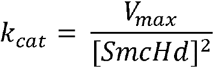

All equations where fit to the experimental data in GraphPad Prism 7. The final protein concentration in the assay was 0.15 μM Smc dimers and 10 μM SmcHd/ScpA-N (unless indicated otherwise) in ATPase assay buffer (50 mM HEPES-KOH pH 7.5, 50 mM NaCl, 2 mM MgCl2, 1 mM NADH), and measurements were carried out at 25 °C.

#### Cysteine Cross-linking of SmcH-CC100

Cross-linking was done as reported in (Soh et al., 2015). SmcH-CC100 protein (2.5 μM dimers) and double-stranded oligonucleotides (40 bp 10 μM) were mixed in 50 mM Tris, 50 mM NaCl, 2 mM MgCl2 and 0.25 mM TCEP (pH 7.5) at room temperature and incubated for 5 min. After incubation, BMOE was added (0.5 mM final). Reactions were incubated for another 5 min at room temperature and quenched with 2-mercaptoethanol (14 mM).

#### Chromatin Immunoprecipitation

The procedure was carried out as described in (Burmann et al., 2017). Cultures of 200 mL SMG were inoculated to OD600 = 0.004 and grown to OD600 = 0.02 at 37°C. Cells were treated with 20 mL of buffer F (50 mM Tris-HCl pH 7.4, 100 mM NaCl, 0.5 mM EGTA pH 8.0, 1 mM EDTA pH 8.0, 10% Formaldehyde) and incubated for 30 min at room temperature with sporadic manual shaking. Cells were harvested by filtration and washed in PBS. The cell mass equivalent to 2 OD units was re-suspended in 1 mL TSEMS (50 mM Tris pH 7.4, 50 mM NaCl, 10 mM EDTA pH 8.0, 0.5 M sucrose and protease inhibitor cocktail (Sigma)) containing 6 mg/mL lysozyme. Protoplasts were prepared by incubating at 37°C for 30 min with shaking. Protoplasts were washed in 2 mL TSEMS, re-suspended in TSEMS, split into 3 aliquots and pelleted. Pellets were flash frozen and stored at −80°C.

Each pellet was re-suspended in 1 mL buffer L (50 mM HEPES-KOH pH 7.5, 140 mM NaCl, 1 mM EDTA pH 8.0, 1% Triton X-100, 0.1% Na-deoxycholate) containing 0.1 mg/mL RNase A and protease inhibitor cocktail. The suspension was sonicated in a Covaris E220 water bath sonicator for 5 min at 4°C, 100 W, 200 cycles, 10 % load and filling level 5. The sonicated material was centrifuged at 4°C and 20,000 × g and 100 μL were separated as input reference. The immunoprecipitation was carried out by mixing 750 μL of the extract with 50 μL of Dynabeads Protein-G freshly charged with 50 μL Anti-ScpB antiserum and incubated for 2 hr on a wheel at 4°C. Beads were washed at room temperature in 1 mL each of buffer L, buffer L5 (buffer μL containing 500 mM NaCl), buffer W (10 mM Tris-HCl pH 8.0, 250 LiCl, 0.5% NP-40, 0.5% Na-Deoxycholate, 1 mM EDTA pH 8.0) and buffer TE (10 mM Tris-HCl pH 8.0, 1 mM EDTA pH 8.0). Beads were resuspended in 520 μL buffer TES (50 mM Tris-HCl pH 8.0, 10 mM EDTA pH 8.0, 1% SDS). The reference sample was mixed with 100 μL buffer L, 300 μL buffer TES and 20 μL 10% SDS. Formaldehyde cross-links were reversed over-night at 65°C with vigorous shaking.

For phenol/chloroform extraction, samples were cooled to room temperature, vigorously mixed with 500 μL phenol equilibrated with buffer (10 mM Tris-HCl pH 8.0, 1 mM EDTA) and centrifuged for 10 min at 20,000 × g. Subsequently, 450 μL of the supernatant was vigorously mixed with 450 μL chloroform and centrifuged for 10 min at 20,000 × g. For DNA precipitation, 400 μL of the supernatant were mixed with 1.2 μL GlycoBlue, 40 μL of 3 M Na-Acetate pH 5.2 and 1 mL ethanol and incubated for 20 min at −20°C. Samples were centrifuged at 4°C and 20,000 × g for 10 min, and the precipitate was washed in 500 μL of 70 % ethanol, dissolved in 250 μL buffer PB (QIAGEN) for 15 min at 55°C, purified with a PCR purification kit (QIAGEN), and eluted in 50 μL buffer EB.

For qPCR, samples were diluted in water (1:10 for IP and 1:1000 for input) and duplicate 10 μL reactions (5 μL master mix, 1 μL of 3 μM primer mix, 4 μL sample) were run in a Rotor-Gene Q machine (QIAGEN) using NoROX SYBR MasterMix (Takyon) and the primer pairs listed in Table S5.

For deep-sequencing, DNA was fragmented to ~200 bp and libraries were prepared using the Ovation Ultralow Library Systems V2 Kit (NuGEN) with 15 PCR cycles. Single-read sequencing was performed on a HiSeq 2500 (Illumina) with 125 bp read length.

#### Analysis of qPCR Data

The threshold cycle (CT) was obtained by analyzing the fluorescence raw data in the Real-Time PCR Miner server (http://ewindup.info/miner/) (Zhao et al., 2005).

IP/input ratios were calculated as α 2^ΔCT^, where ΔCT = CT (Input) – CT (Alipour and Marko) and α is a constant determined by extraction volumes and sample dilutions. Data are presented as the mean of duplicates.

#### Analysis of ChIP-seq Data

Data was analysed as described in (Bürmann et al., 2017). Deep-sequencing data for the immunoprecipitate were mapped to the B. subtilis reference genome NC_000964 (centered on its first coordinate) using Bowtie for Illumina in the Galaxy project website (https://usegalaxy.org/). Reads were filtered for mapping quality (MAPQ) greater than 10, reduced to bins of 250 bp, and normalized for total read count in SeqMonk. Data are presented in reads per million (rpm).

For ratiometric analysis, the reduced data of each sample was compared to the reduced data of the wild-type sample (Minnen et al., 2016). For each bin, the larger value was divided by the smaller, and the resulting ratio was plotted above the coordinate axis for value (mutant) ≥ value (WT) and below the axis otherwise. Data was exported and graphically represented in GraphPad Prism 7 for Mac.

#### In vivo cross-linking

The procedure was based on the one described in (Bürmann et al., 2013). Cultures of 200 mL SMG were grown to exponential phase (OD600 = 0.02) at 37°C. Cells were harvested by filtration, washed in cold PBS + 0.1% glycerol (PBSG), and split into aliquots of a biomass equivalent to 0.85 OD units. Cells were centrifuged 2 min at 10,000 *g,* re-suspended in 200 μL PBSG and cross-linked by adding 0.5 mM BMOE. Cells were incubated with BMOE for 10 min on ice. The reaction was quenched by the addition of 14 mM 2-mercaptoethanol. Cells were pelleted and resuspended in 30 μL of PBSG containing 75 U/mL ReadyLyse Lysozyme, 750 U/mL Sm DNase, 5 μM HaloTag TMR Substrate and protease inhibitor cocktail (Sigma). Lysis was performed at 37°C for 15 min. After lysis, the material was diluted with 10 μL of 4X LDS-PAGE buffer, samples were incubated for 5 min at 95°C and resolved by SDS-PAGE. Gels were imaged on a Typhoon FLA9000 (GE Healthcare) with Cy3 DIGE filter setup.

#### Chromosome Entrapment Assay

Agarose plug assays were done as described in (Wilhelm and Gruber., 2017). The protocol was adopted for use of agarose microbeads instead of plugs following procedures described in (Wing et al., 1993). The equivalent of 3.75 ml OD cell mass was resuspended in 121 μL of PBSG, crosslinked in the presence of 1 mM BMOE and quenched with 28 mM 2-mercaptoethanol. Each sample was split into two aliquots, one of 45 μL labelled as input and the other of 90 μL labelled as beds. The bead sample was mixed with 9 μL of DynaBeads (Invitrogen) (to label the otherwise translucent microbeads) and 1x PIC (Sigma). The samples were prepared one at a time: first, the cell suspension was incubated for 10 s at 45 °C and immediately mixed with 100 μL of liquid 2 % low-melt agarose (BioRad) prewarmed to 45 °C. 800 μL of mineral oil at 45 °C (sigma) was added and vigorously mixed for 1-2 minutes at 4 °C. The mixture was kept at room temperature until all samples were prepared. The oil was removed by centrifugation for 1.5 min at 10,000 g at room temperature. The agarose microbeads were rinsed twice in 1 mL of room temperature PBSG. Cell lysis was achived by addition of ReadyLyse Lysozyme (4 u/μL final), protease inhibitor, EDTA pH 8 (1 mM final) and Halo-TMR substrate (5 μM final) and incubation for 25 min at 37 °C. Separate input samples were lysed in the presence of Sm DNase (5 u/μL final) without EDTA. The input samples where then mixed with 50 μL of 2X LDS loading buffer and stored at -20 °C until used.

The agarose beads were then washed twice in PBSG at room temperature supplemented with 1 mM EDTA pH 8 and then incubated twice in 1 mL of TES buffer (50 mM Tris-HCl pH 8.0, 10 mM EDTA pH 8.0, 1% SDS) at room temperature on a rotating wheel. The final wash was done over-night at 4 °C in TES buffer on a rotating wheel. SDS and EDTA were removed by rinsing the beads three times with 1 mL PBS. Microbeads were resuspended in 100 μL of PBS and incubated with 50 units of Sm DNase 30 min at 37 °C protected from light. The mixture was incubated at 70 °C for 1 min with vigorous shaking, cooled for 5 min at 4 °C and centrifuged at 21,000 rpm for 15 min at 4 °C. The supernatant was transferred to a 0.45 μm Costar Spin-X Tube Filter (Corning) and spun for 1 min at 10,000 × g. The flow-through was diluted to 1 mL final volume with PBS and proteins were precipitated by 10 % (w/v) TCA in presence of 3 μg of BSA (Koontz, 2014). The precipitate was re-suspended in LDS Sample Buffer (NuPage) and heated for 5 min at 95 °C. Samples were loaded on a 3%–8% Tris-Acetate gel (Life Technologies) and run for 1.5 hr at 35 mA per gel at 4°C aside with 10 % of the input material. Gels were scanned on a Typhoon scanner (FLA 9000, GE Healthcare) with Cy2-DIGE (for Oregon-Green) or Cy3-DIGE (for TMR) filter setting.

## Supporting information

## Data availability

ChIP-Seq data reported in this manuscript will be deposited at the NCBI Sequence Read Archive. Original data will be made available at Mendeley data.

## Author contributions

L.B.R.A., project development and initial experiments on DNA entrapment in S and K compartments; R.V.N., optimization and application of chromosome entrapment assay using agarose microbeads; R.V.N., all other experiments; R.V.N., L.B.R.A., S.G., conception of experiments and interpretations of data; R.V.N., S.G., preparation of manuscript.

## Acknowledgements

We are grateful to all members of the Gruber lab for critical comments on the manuscript and to Kim Nasmyth and Christophe Chapard for sharing unpublished results. We thank Yolanda Schaerli for advice on the fabrication of agarose microbeads and Anna Anchimiuk for providing genetic tools. Deep sequencing was performed at the Genome Technology Facility (GTF) at UNIL. This work was supported by a European Research Council Consolidator Grant (‘Chrocodyle’ #724482 to S.G.) and the University of Lausanne.

## Supplemental Figure Legends

**Figure S1. Related to Figure 1.**

(A) Size exclusion chromatography of SmcH-CC100 proteins. Gel filtration profile on a Superdex 200 10/300 column.

(B) Dilution spotting of *Smc(3E-hng)* and *Smc(3A-hng)* cells on nutrient-rich medium (ONA) grown for 12 hours (upper panel) or 16 hours (lower panel). The lower panel is identical to the one shown in Figure 1E.

**Figure S2. Related to Figure 2**.

(A) Dilution spotting of single glutamate mutants. R57 is buried in E heads Smc and contacts ATP, thus it was not considered to be a DNA binding residue. As in Figure 1E.

(B) Dilution spotting of five single alanine mutants. As in (A).

(C) Dilution spotting of all double alanine mutant combinations. As in (A).

(E) Top view of *G. stearothermophilus* SmcHd-ATPγS structure (PDB: 5H68) in gray, superimposed with the Rad50Hd-ATPγS-DNA structure (PDB: 5DNY). Only DNA is shown in yellow colors. In blue are shown the five positively charged residues identified in Figure 2C and D. ATPγS not shown. As in Figure 2D using another Rad50-ATPγS-DNA co-structure for superimposition. Top view (left panel) and side view (right panel).

**Figure S3. Related to Figure 3.**

(A) Chromatogram of SmcHd/ScpA^N^ elution in a Superdex 200 10/300 size exclusion column. As experiments as in Figure 3C showing wider range.

(B) DNA binding curves represented by the fluorescence anisotropy (linear scale) change at low protein concentrations. Same data as in Figure 3D.

**Figure S4. Related to Figure 4.**

(A) Specific ATP hydrolysis activity of SmcHd/ScpA^N^ mutants (axis and data in blue colors) and full-length Smc (axis and data in black colors) at increasing protein concentrations. Each rate was divided by the appropriate protein concentration and fit to the first order *k_cat_* equation for full-length Smc dimer, or to the second order *k_cat_* equation for the SmcHd/ScpA^N^ variants (see Experimental procedures).

(B) ATP hydrolysis rate of the full-length Smc(5A-hd) protein. As in Figure 4C.

**Figure S5. Related to Figure 5.**

(A) Chromosome-wide ChIP-seq profiles (in reads per million) for data shown in Figure 5B.

(B) Ratiometric analysis of chromosome-wide profiles shown in (A). Ratios were calculated as described in Figure 5C.

(C) ChIP-qPCR against ScpB for Smc(5A-hd). As in Figure 5D.

(D) Schematic overview of cross-linking intermediates taken from (Wilhelm et al., 2015).

**Figure S6. Related to Figure 6**.

(A) Dilution spotting of *Smc(6A)* and *Smc(5A-hd, 3A-hng)* strains. Growth on ONA was monitored at 12 and 16 hours (upper and middle panels).

(B) Schematic view of K1151C cross-linking of E heads but not J heads. As in Figure 6C.

**Figure S7. Related to Figure 7.**

(A) Scheme of the chromosome entrapment assay using agarose microbeads instead of agarose plugs. Cells are incubated with BMOE and mixed with melted agarose solution and with mineral oil. An emulsion is formed by continuous vortexing until the agarose forms a gel. The oil is removed and the cells inside the agarose microbeads are lysed and washed in presence of SDS. The proteins retained in the agarose microbeads are released by DNA digestion and precipitated to finally resolve them in an SDS-PAGE and detect them by in-gel fluorescence.

(B) Chromosome entrapment assay using agarose plugs for strains harboring the Smc arm cross-linking residue D280C. As in Figure 7B.

