## Supplementary figures and images for "Separate Compartments for Chromosome Entrapment and DNA Binding during SMC translocation"

### Supplementary file 1

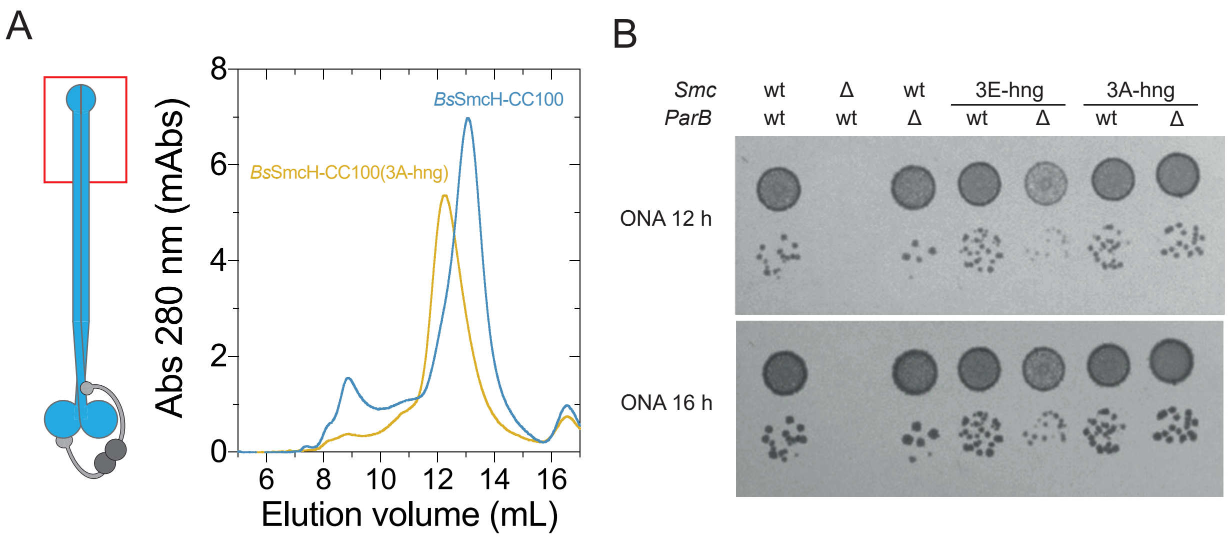

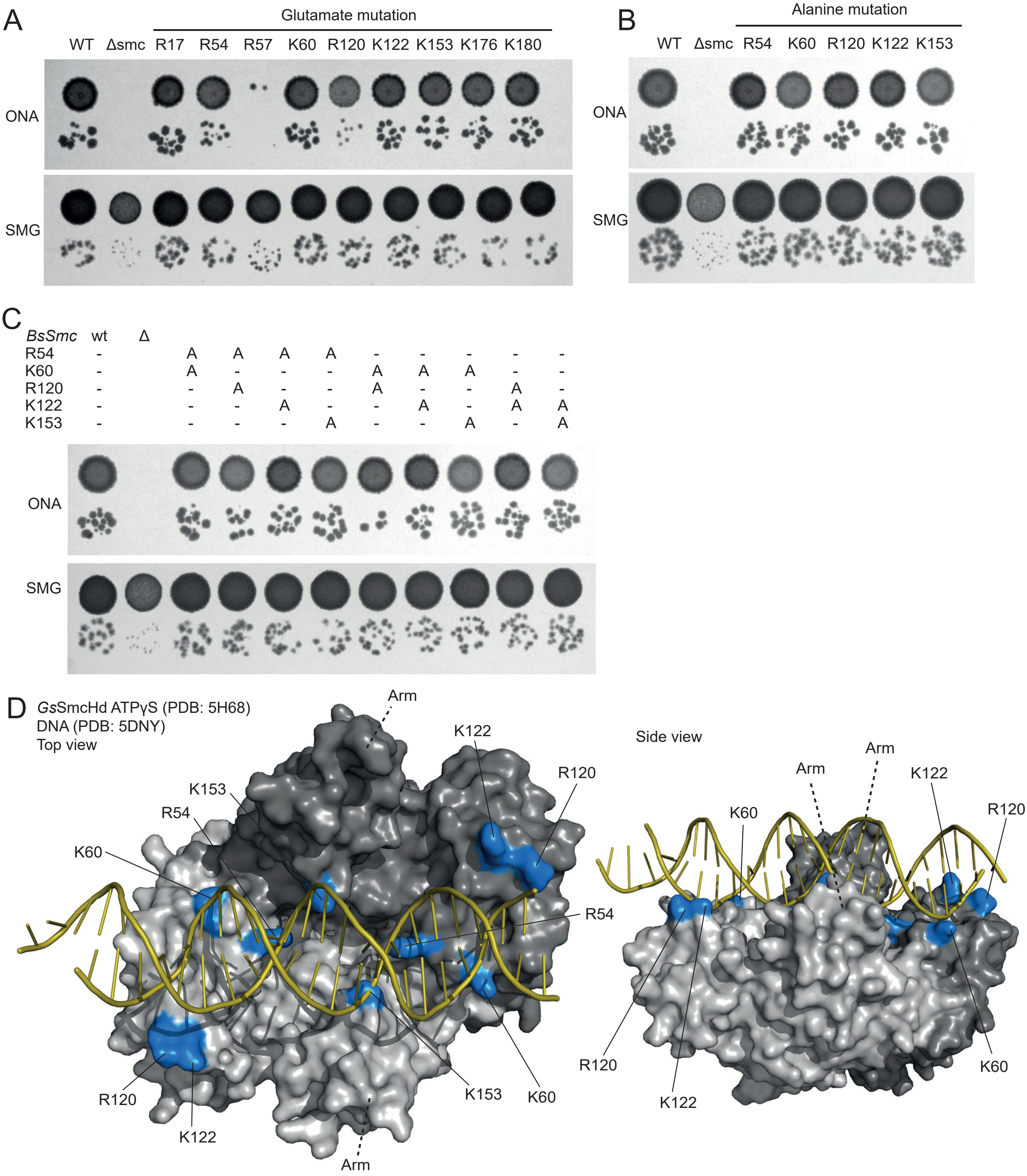

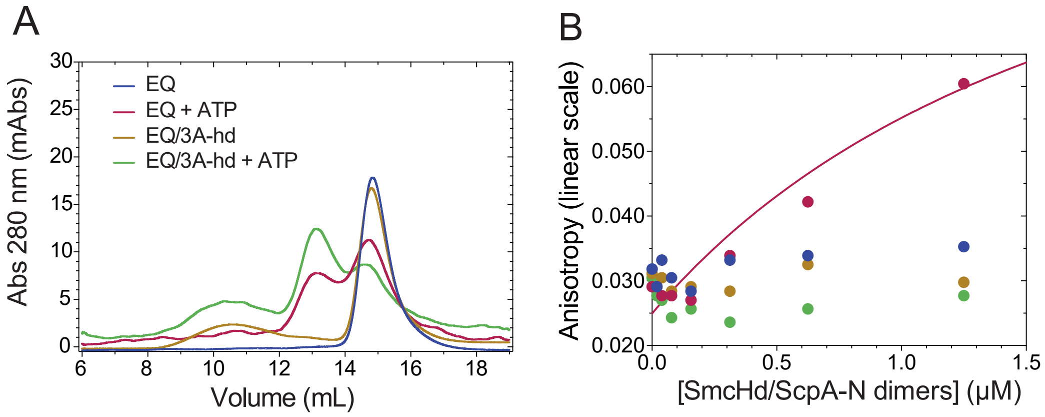

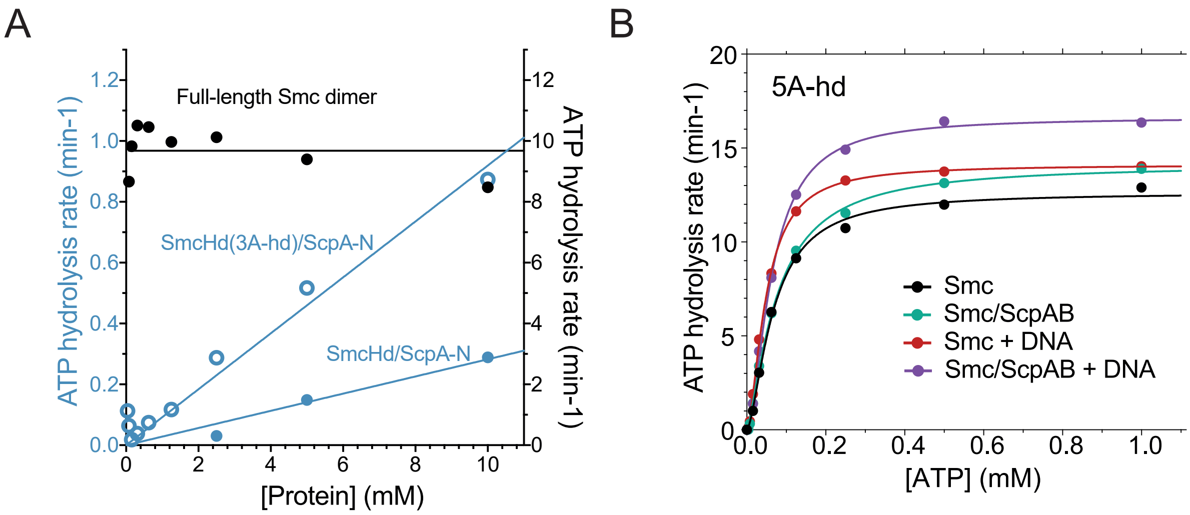

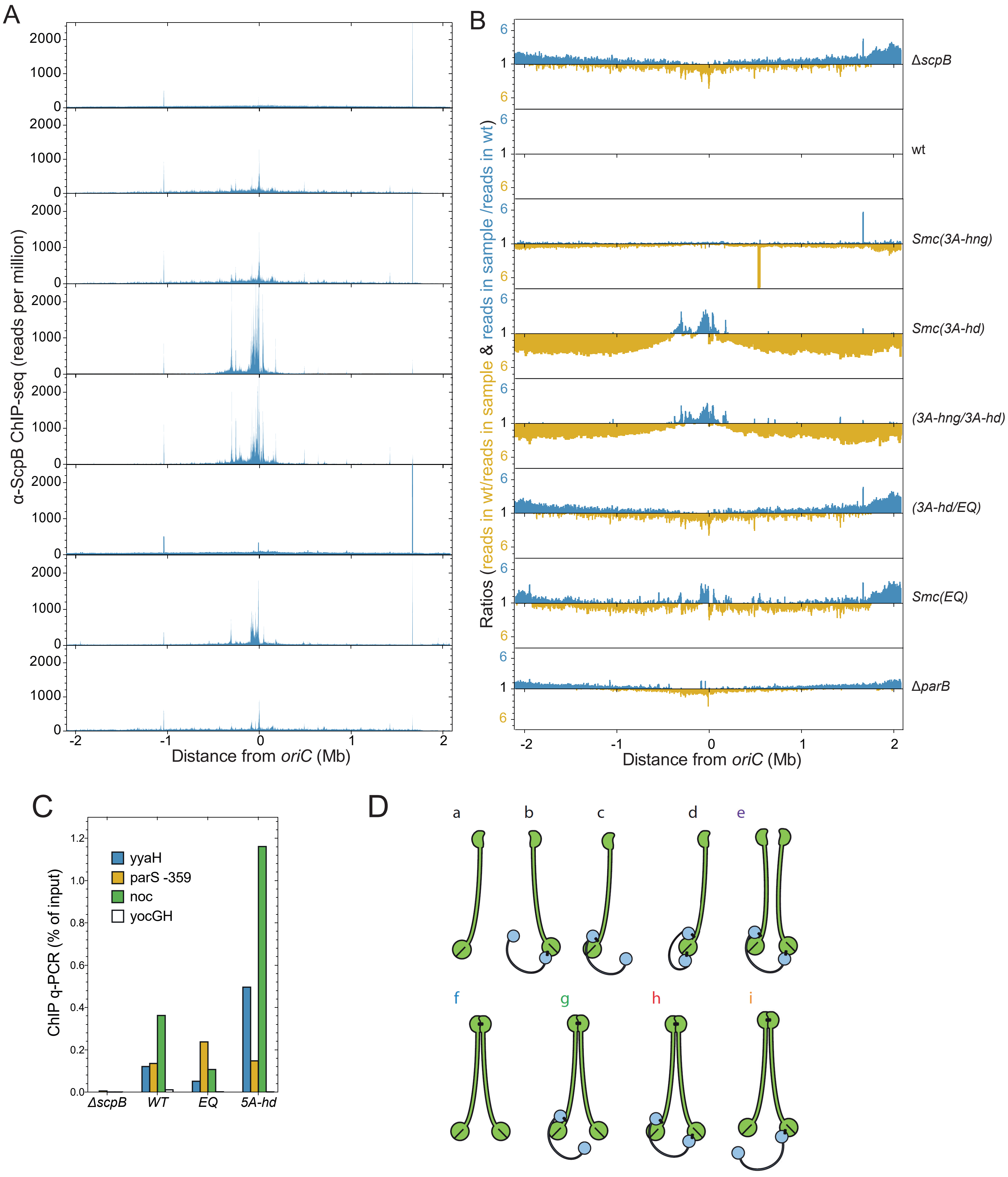

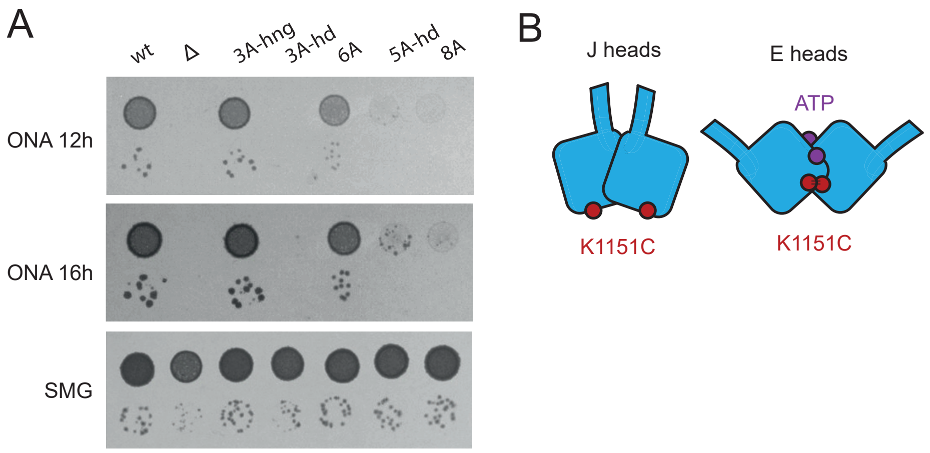

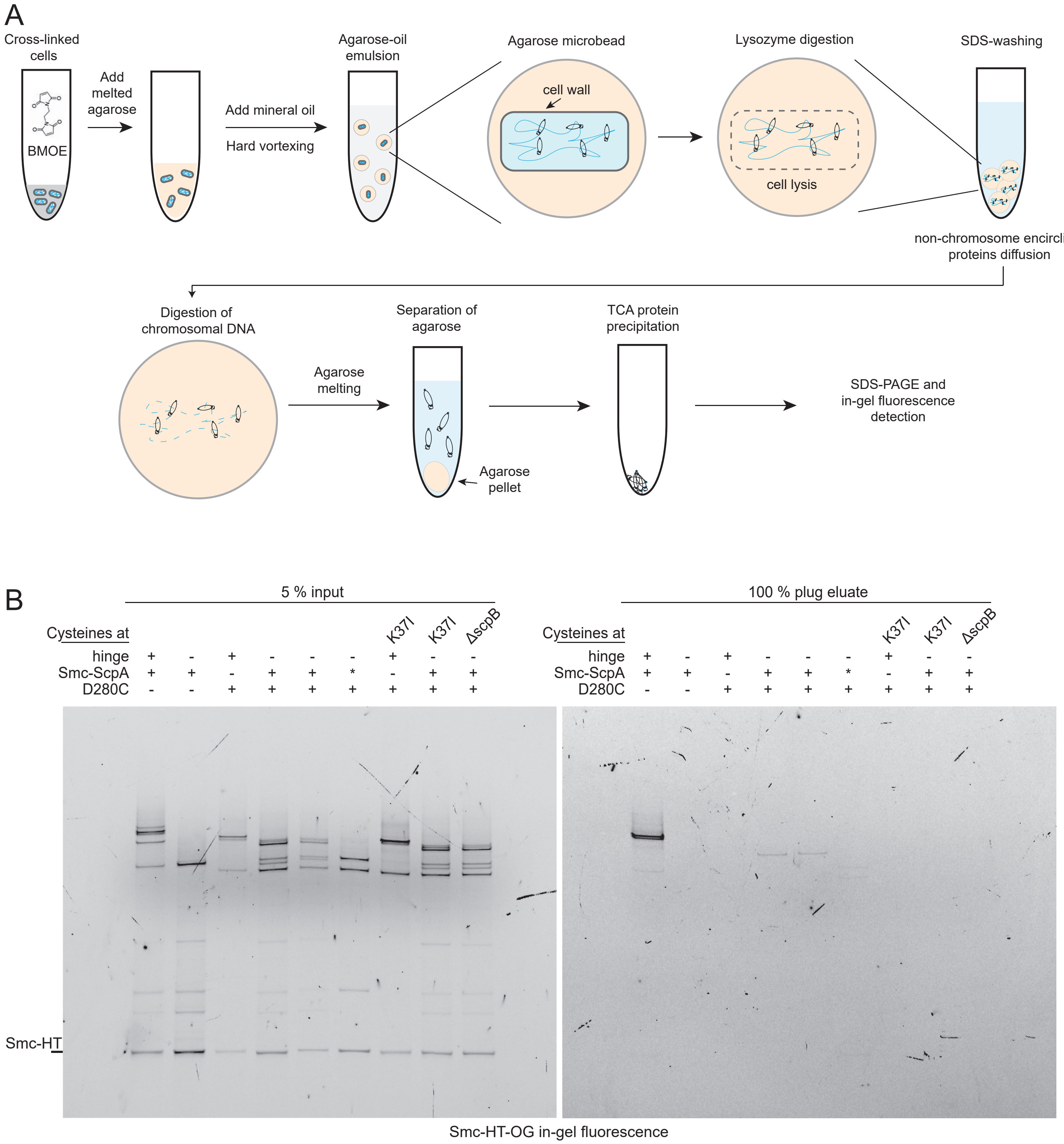
